# ARX regulates cortical interneuron differentiation and migration

**DOI:** 10.1101/2024.01.31.578282

**Authors:** Youngshin Lim, Shyam K Akula, Abigail K Myers, Connie Chen, Katherine A Rafael, Christopher A Walsh, Jeffrey A Golden, Ginam Cho

## Abstract

Mutations in aristaless-related homeobox (*ARX*) are associated with neurodevelopmental disorders including developmental epilepsies, intellectual disabilities, and autism spectrum disorders, with or without brain malformations. Aspects of these disorders have been linked to abnormal cortical interneuron (cIN) development and function. To further understand ARX’s role in cIN development, multiple *Arx* mutant mouse lines were interrogated. We found that ARX is critical for controlling cIN numbers and distribution, especially, in the developing marginal zone (MZ). Single cell transcriptomics and ChIP-seq, combined with functional studies, revealed ARX directly or indirectly regulates genes involved in proliferation and the cell cycle (e.g., *Bub3*, *Cspr3*), fate specification (e.g., *Nkx2.1*, *Maf*, *Mef2c*), and migration (e.g., *Nkx2.1*, *Lmo1*, *Cxcr4*, *Nrg1*, *ErbB4*). Our data suggest that the MZ stream defects primarily result from disordered cell-cell communication. Together our findings provide new insights into the mechanisms underlying cIN development and migration and how they are disrupted in several disorders.

## Introduction

The mammalian cerebral cortex primarily consists of excitatory projection neurons (PNs) and inhibitory interneurons (INs) ^1–4^. Glutamatergic PNs mediate communication between different brain regions by connecting with local networks as well as with distant targets ^5^. GABAergic INs modulate neural circuits through local connections with PNs and other INs ^4^. Perturbations in the balance between neuronal excitation and inhibition are apparent in numerous neurological and neuropsychiatric disorders, including subtypes of autism spectrum disorder, many developmental epilepsies, and schizophrenia^6–10^. Thus, understanding the mechanisms that control the differentiation, migration, and integration of cortical INs (cINs) will contribute to the development of targeted therapies for these disorders.

cINs have been classified according to their developmental origin, morphology, gene expression, and electrophysiologic firing properties. cINs expressing the calcium binding protein parvalbumin (PV) or the neuropeptide somatostatin (SST) primarily originate from the medial ganglionic eminence (MGE), constitute ∼70% of all cINs in the mature mouse cortex, and populate all cortical layers ^3,4,11^. In contrast, cINs expressing the serotonin receptor 5HTR3a originate from the caudal ganglionic eminence (CGE) account for ∼30% of all cINs and preferentially populate upper cortical layers ^3,4,11^. Those derived from the preoptic area are a small portion of cINs ^1,2,4,12^. These major IN groups can be further divided into subclasses based on features such as synaptic specificity and gene expression ^4^. Recent advances in single cell transcriptomics have identified more than 50 transcriptionally defined subtypes of cINs using unbiased clustering methods^13^. How these diverse groups of cINs are generated and differentiate into different subtypes as well as how their migration to the cortex is regulated remain incompletely understood.

The spatial and temporal expression of multiple transcription factors across the ganglionic eminences (GEs) play key roles in the specification of cIN subtypes. For example, the transcription factor NKX2.1 is essential for the specification of SST and PV INs in the MGE^12,14^. Moreover, after its initial expression in the progenitor cells of the MGE, subsequent downregulation of *Nkx2.1* is required for the migration of MGE-derived INs toward the cortex^15^. It has also been shown that MAF and MAF-b (both transcription factors) modulate MGE-derived IN differentiation by suppressing SST cell identity and promoting a PV fate^16,17^. Overall, the establishment of IN subtypes is a complex process that involves the coordinated spatial and temporal expression of multiple transcription factors in response to extracellular signaling molecules^11,18–20^.

In addition to the diversity of cINs, their complex path of migration is another important feature of cIN development^3,11,21,22^. cINs are generated in the ventral telencephalon including MGE, CGE, and preoptic area (although a minor subset appears to originate in the dorsal telencephalon in humans ^23–26^), from where they migrate along defined paths to enter the developing cortex. The initiation of IN migration involves chemorepulsive signals from the proliferative zones of the GEs ^27^ and motogenic factors ^28–31^. Cortically directed INs use NRP1/SEMA3A signaling ^32^ to navigate around the developing striatum and migrate to the cortex using chemoattractant guidance cues such as CXCL12 and NRG1^33–35^. Once they cross from the ventral to the dorsal telencephalon, cINs migrate along different streams: the most superficial within the marginal zone (MZ), a second deep to the developing cortex within the intermediate zone (IZ)/subventricular zone (SVZ), and a third developing later in the subplate (SP) (E15-16 in the mouse) ^36,37^. Interestingly, the choice of migration stream seems to be coupled to axon targeting programs to optimize the inhibitory circuit assembly ^38^. The final step is a timed exit from the migratory stream and invasion into the cortical plate via radial migration, a process also modulated in part by CXCR4/CXCL12 signaling ^39^.

*ARX* is a transcription factor encoded on the X-chromosome that is expressed in cortical PN and IN progenitor cells. Its expression persists in postmitotic INs but not in cortical PNs^40^. We have interrogated a series of *Arx* mutant mouse models to understand the relationship between the ARX transcriptional networks in IN development and the phenotypic consequences related to human disorders^41^. These models include conditional *Arx* knock-out mice (*Arx* cKO) generated by crossing *Arx* floxed mice^42^ with four different IN-targeted *Cre* lines (*Gad2Cre*, *Nkx2.1Cre* or *Nkx2.1CreER*, *SstCre*, and *Dlx5/6Gre*), and an alanine tract expansion mouse model (*Arx^(GCG)7^*)^41^. Single cell transcriptomics and ARX ChIP-seq revealed the role of ARX and its downstream target genes in multiple aspects of cIN development: i) transition from proliferation to differentiation, ii) SST+ and PV+ IN subtype specification, and iii) IN migration, particularly exiting the GE and initial entry into the cortical MZ stream. Though less severe, *Arx^(GCG)7^* brains exhibit similar though not identical defects. Together, these findings provide new insights into the mechanisms underlying cIN development and ultimately how different mutations can explain the spectrum of overlapping phenotypes found in patients.

## Results

### Loss of Arx in the GE results in abnormal numbers and distribution of cINs

To study the role of ARX in IN development, we crossed *Arx^flox/+^;Ai14/Ai14* female mice (*Ai14* is a CRE-dependent tdTomato reporter) ^42,43^ with *Gad2Cre* males [expressing CRE in the *Gad2* (glutamate decarboxylase2, pan-IN marker)-lineage beginning on E10 in the GE SVZ] ^44^ (Fig. 1a). The resulting *Arx* cKO mice (*Arx^flox/y^*; *Ai14/+*; *Gad2Cre/+*) die shortly after birth, similar to constitutive knockout (KO) mice^41^. The overall brain size and morphology of the *Gad2Cre Arx* cKOs appeared normal at E14.5 (data not shown).

**Figure 1.**
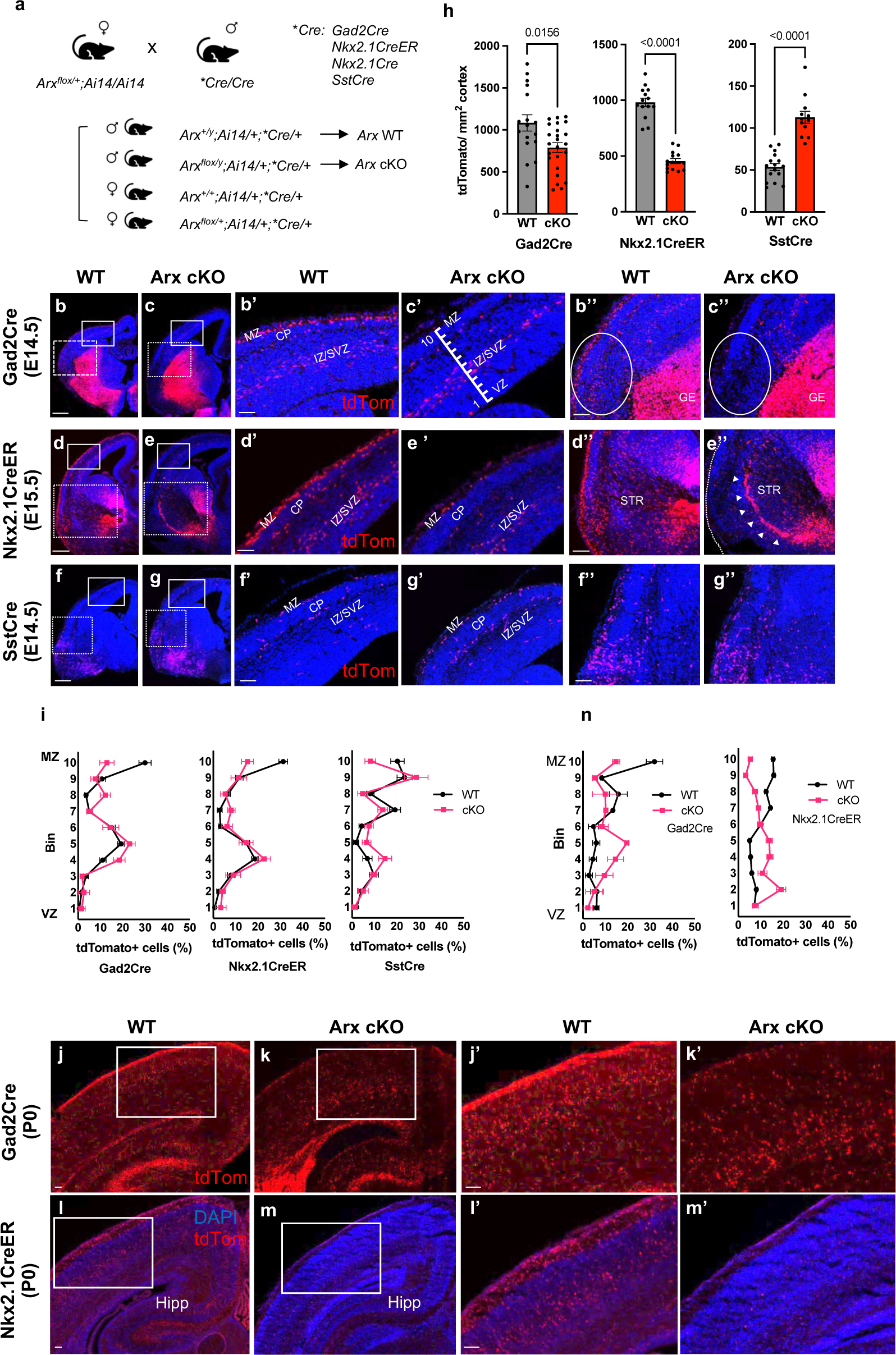
Cortical interneuron (cIN) density and distribution defects in *Arx* cKO;Ai14 (simply referred to as cKO) mice driven by *Gad2Cre*, *Nkx2.1CreER*, and *SstCre*. **a** Overview of the mouse lines used and breeding strategy. **b-g, j-m** Representative images of the immunolabeling of tdTomato (red) on coronal sections of WT and *Arx* cKO under the *Gad2Cre* (**b**, **c, j, k**), *Nkx2.1CreER* (**d**, **e, l, m**), and *SstCre* (**f**, **g**) taken at E14. 5 (**b**-**g**) or P0 (**j-n**). DAPI (blue) was used for nuclear staining. b’-g’ and j’-m’ magnified images of the lined boxed areas in b-g and j-m, respectively; b’’-g’’ are magnified images of the dotted boxed areas in b-g. Ten equally distributed bins are indicated in c’; circled areas in b’’-c’’ indicate examples of the cortical entry area of the INs; white arrowheads in e’’ indicate accumulated clusters of tdTomato+ INs at the striatal boundary; dotted line in e’’ denotes the boundary of the section. **h** Quantification of tdTomato+ cell density in WT and cKO neocortices from each indicated *Cre* driven mice at E14.5. n = 17, 24, 14, 14, 16, 12 sections from at least 4 hemispheres/genotype for each bar graph (left to right). **i, n** Quantification of the percentage of the tdTomato+ cells in each bin to the total tdTomato+ cell numbers in all 10 bins from each indicated Cre driven mice at E14.5 (i) or P0 (n). n = 16, 16, 13, 14, 16, 9 sections from at least 4 hemispheres/genotype for each graph (left to right; WT to cKO order). CP, cortical plate; GE, ganglionic eminence; Hipp, hippocampus; IZ, intermediate zone; MZ, marginal zone; SVZ/VZ, subventricular zone/ventricular zone; STR, striatum. Scale bars in b, d, f: 400 μm; b’, b’’, d’, d’’, f’’, j, j’, l, l’: 100 μm. Error bars in each graph: mean ± sem. Numbers over the bars: p values.

The total number of tdTomato+ cINs was reduced (Fig. 1b, b’, c, c’, h and Supplementary Fig. 1) predominately due to reductions in the MZ migration path (Fig. 1b’, c’, i). There was a near complete absence of cINs in the ventral most aspect of the GE-cortical boundary (compare circled areas in Fig. 1b’’, c’’) where the cortical MZ migration stream starts. Consistent with E14.5 results, at PO, there were dramatically fewer cINs in cKO mice compared to WT, and tdTomato^+^ cINs in the cortex were disproportionally localized in deeper cortical layers and few in the upper layers (Fig. 1j, k, j’, k’, n).

Given that *Gad2-Cre* is expressed in all GEs (MGE/LGE/CGE), we next sought to investigate specific roles of ARX in MGE-derived cINs using an *Nkx2.1-CreER* line (*Nkx2.1* is expressed in the MGE and not the CGE or LGE) ^44^. *Nkx2.1CreER-Arx cKO* mice were generated similarly to *Gad2Cre-Arx cKO* mice (Fig. 1a); tamoxifen was injected at E12.5 to induce CRE-mediated *Arx* deletion. These cKO mice showed similar IN phenotypes described above, with a dramatic reduction of MZ cINs and relative sparing of those in the IZ/SVZ (Fig. 1d, d’, e, e’, h, i and Supplementary Fig. 2a). Remarkably, many cINs migrating out of the MGE appeared stuck around the developing striatum, failing to migrate to the cortex (arrowheads in Fig. 1e’’), compared to the broadly distributed cINs in WT (Fig. 1d’’). Similarly, by P0, most of the *Arx*-deficient *Nkx2.1*-lineage (tdTomato+) INs failed to migrate into the cortex and hippocampus (Figure 1l, l’, m, m’, o). These findings were also observed in *Nkx2.1Cre*-driven cKO mice (tamoxifen independent) (Supplementary Fig. 2b, c). Our data suggest that one role for ARX in MGE-derived cells is to enable entry to the cortical MZ stream.

We next explored the consequences of ARX depletion in specific IN subpopulations using the *Sst-Cre* which is expressed in postmitotic SST neurons soon after they exit the MGE SVZ ^44^. At E14.5, contrary to the prior cKO models, the cortical entry of the *SstCre-Arx* cKO tdTomato+ INs into the MZ stream from the GE did not appear to be severely reduced relative to WT (Fig. 1f, f’, f’’ g, g’, g’’ and Supplementary Fig. 3), although the proportion of cKO tdTomato+ cells in the MZ stream (over the total tdTomato+ cells) is still lower than that of the WT littermate controls (Fig. 1i). Curiously, the cortical density of tdTomato+ INs was higher in the *Sst-Cre Arx* cKOs relative to WTs (Fig. 1h). These results suggest that ARX’s function in SST+ INs is required for their distribution in the cortical MZ migration stream. Taken together our data across multiple *Arx* cKO lines indicate ARX plays important roles in generation of cINs and their migration in the MZ pathway.

### Arx is necessary for GE progenitor cells to exit the cell cycle and for cINs to enter the cortex

We focused on *Gad2Cre*-*Arx* cKO mice to characterize the role *Arx* plays in proliferation, differentiation, and migration of cIN precursors. We first examined cell proliferation rate by analyzing a 1.5 hr incubation period after EdU was injected at E13.5. The proportion of tdTomato+ cells labeled with EdU was smaller (though not reaching statistical significance) in the *Gad2Cre*-*Arx* cKO mice (0.61 ± 0.09 and 0.75 ± 0.05; p = 0.1758), suggesting that the loss of ARX might have minimal impact, if any, on the GE cell proliferation rate (Fig. 2a, b).

**Figure 2.**
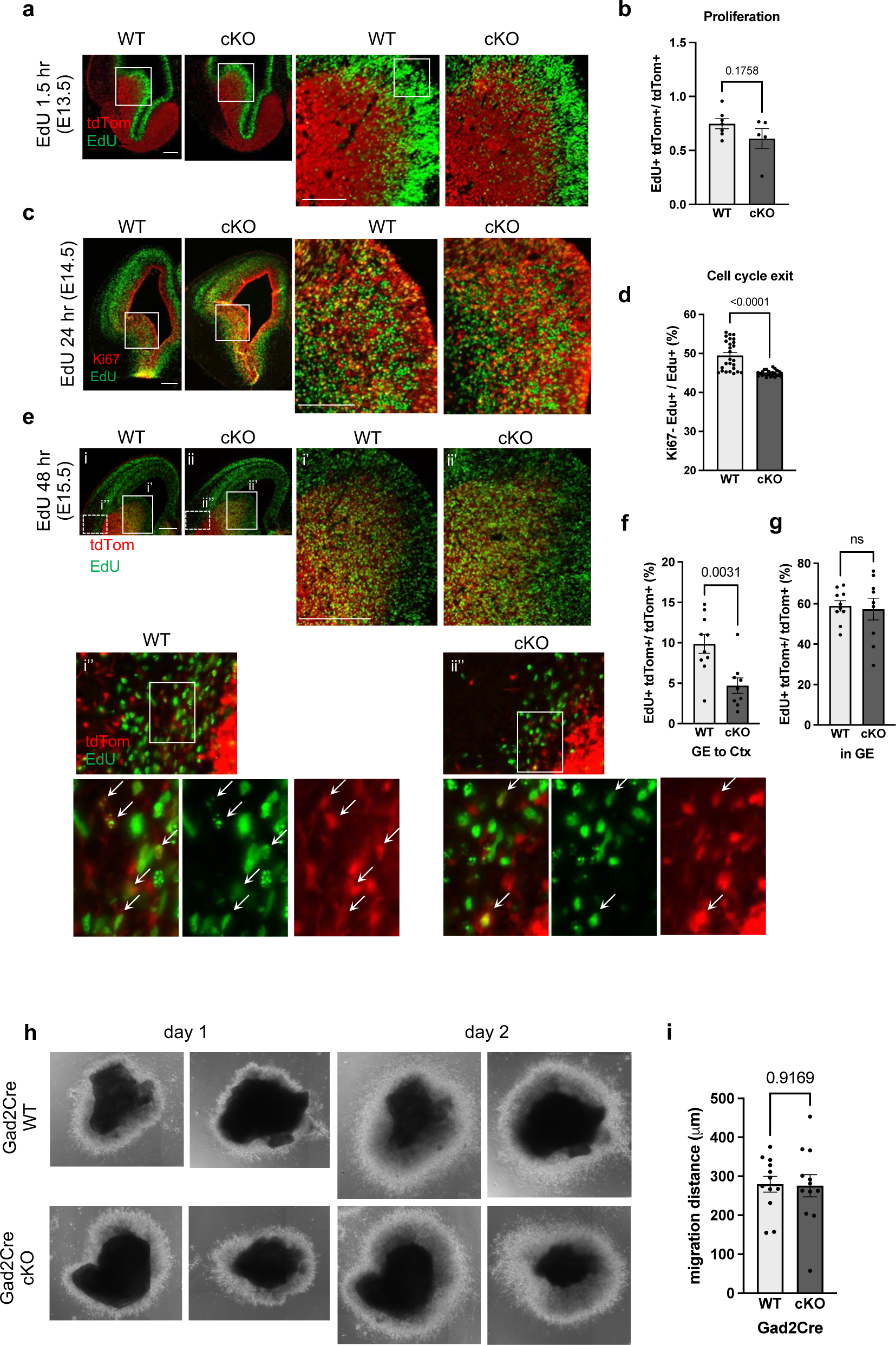
ARX depletion leads to reduced cell cycle exit and cortical entry of INs. **a** Representative images of EdU (green) and tdTomato (red) immunolabeling on coronal sections (GE) of the WT and cKO (*Gad2Cre*) embryonic brains with EdU injected at E13.5 and incubated for 1.5 hr. Right panels are magnified images from the boxed areas in the left panels. **b** Quantification of the cell cycle index (EdU, tdTomato double positive cells/tdTomato+ cells) from the images taken as in a. n = 7 (WT), 5 (cKO) sections from 4 hemispheres. **c** Representative images of EdU (green) and Ki67 (red) immunolabeling on coronal sections (GE) of the WT and cKO (*Gad2Cre*) embryonic brains with EdU injected at E13.5 and incubated for 24 hr. Right panels are magnified images from the boxed areas in the left panels. **d** Quantification of the percentage of cells exited the cell cycle from the images taken as in c. n = 26 sections from 6 hemispheres. **e** Representative images of EdU (green) and tdTomato (red) immunolabeling on coronal sections (GE) of the WT (i) and cKO (ii) (Gad2Cre) embryonic brains with EdU injected at E13.5 and incubated for 48 hr. i’ and ii’ are magnified images of the lined boxed areas in i and ii; i’’ and ii’’ are magnified images of the dotted boxed areas in i and ii. Each three images below i’’ and ii’’ are magnified images of the boxed areas in i’’ and ii’’. Arrows in the bottom images indicate examples of cells double positive for EdU and tdTomato. **f**, **g** Quantification of the percentage of EdU+ cells among tdTomato+ cells entering to cortex (f) or staying in GE (g). n = 10 (WT), 9 (cKO) sections from 4 hemispheres. **h** Two sets of the representative images of the explant cultures prepared at E13.5 and incubated for 1 day (left panels) or 2 days (right two panels). **i** Quantification of migration distance measured from the explant culture images as shown in h. n = 12 explants from 4 hemispheres. Scale bars: 200 μm. Error bars in each graph: mean ± sem. Numbers over the bars: p values.

Next, we monitored the cell cycle exit rate in the GEs after 24-hour labeling of EdU (injected at E13.5) together with Ki67 immunolabeling of cycling cells (Fig. 2c, d). Unexpectedly, the percentage of cells that exited the cell cycle (EdU+ Ki67−/EdU+) was lower in *Gad2Cre-Arx* cKO GE than in WT (44.91 ± 0.14% and 49.5 ± 0.75%; p < 0.0001), indicating more GE cells in the cKO mice were in cycle and fewer cells exited the cell cycle to differentiate. These data suggest that ARX normally promotes cell cycle exit of the progenitor cells in the GE.

We then tested whether IN migration from the GEs into the cortex was affected. The position of EdU/tdTomato+ labeled INs 48 hrs post-EdU injection at E13.5 showed that a significantly lower proportion of EdU-labeled *Gad2*-lineage INs crossed the boundary between the GE and cortex in cKO mice compared to WT brains (4.7 ± 0.96% and 9.9 ± 1.15%; p = 0.0031) (Fig. 2e, f). In contrast, the numbers of EdU+ *Gad2*-lineage INs located within the GE were comparable in both mice (57.38 ± 5.38% for cKO and 58.95 ± 2.53% for WT; p = 0.7959) (Fig. 2g). These results indicate that the loss of ARX results in a failure of cINs to enter the cortex. To ensure this was not a primary defect in the ability of INs to migrate, a GE explant culture was deployed ^56^ and found no significant difference in migration distance between WT and cKO INs (Fig. 2h-j). Taken together, our data indicate that the reduced numbers of cIN along the MZ path, and eventually in the upper layer of the mature cortex, likely results from a combined defect in the generation of cINs from progenitor cells and a failure of cINs to enter the MZ migration path.

### Cell type and gene regulatory network defects upon loss of Arx

To understand how changes in gene expression contribute to cIN phenotypes, single cell transcriptomic profiling was performed; tdTomato^+^ *Gad2*-lineage cells were isolated from E16.5 WT and cKO (*Gad2-Cre*) brains via fluorescence-activated cell sorting (FACS) and processed for scRNA-seq (Fig. 3a). After quality control, removing doublets, and normalization of sequencing results, we obtained a total of 27,585 high-quality cells from WT and cKO brains (11,224 and 16,361 cells, respectively). UMAP analysis identified several distinct clusters of cells we referred to as *MGE*, *CGE*, *LGE*, *septal,* and *excitatory neuron* clusters based on their marker gene expression ^59,60^. Three *LGE* [*D1 MSN* (medium spiny neuron) *Tshz1* (Teashirt family zinc finger 1), *D1 MSN Pdyn* (Prodynorphin), and *D2 MSN*] clusters were further identified along with two *unknown* clusters (Fig. 3b, d) ^59,60^. The identified excitatory neuron cluster included *tdTomato* transcripts but not *Gad2* (Fig. 3b and data not shown), potentially representing contamination or the loss of *Gad2* expression.

**Figure 3.**
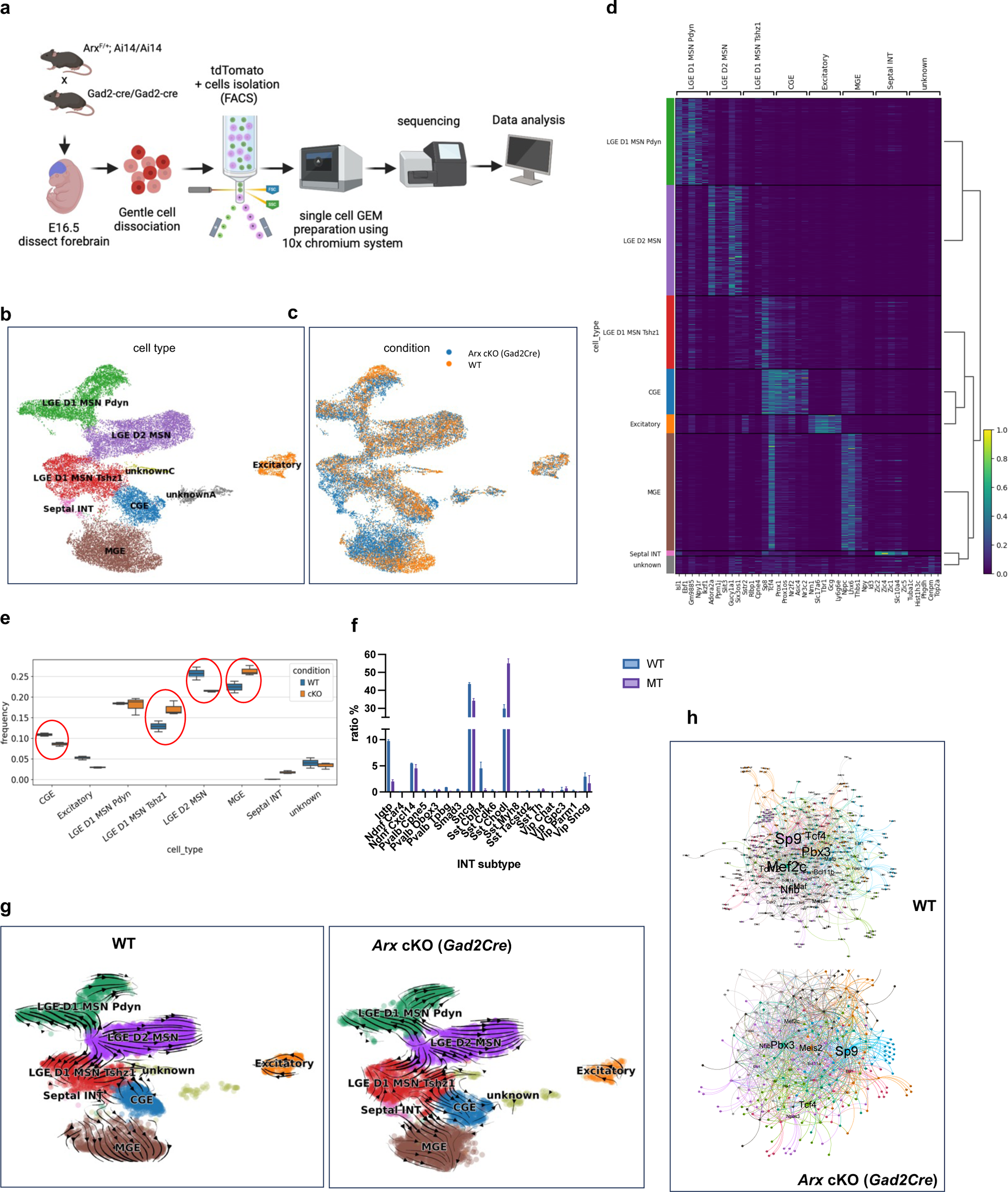
Single cell transcriptome analysis of *Gad2Cre*-mediated *Arx* cKO and WT INs. **a** Schematic diagram of overall experimental approach. The tdTomato+ INs were collected using FACS from E16.5 forebrains (n=3 for WT and n=2 for MT). **b-c** UMAP projection of INs identified 8 clusters (b) and the results from WT (orange) and cKO (blue) were overlaid to each other (c). **d** Classification of INs into several IN subtypes. Heatmap of clusters indicates expression of maker genes differentially expressed that best present interneuron subtypes. **e** Composition analysis of clusters. Red circles are examples of clusters showing differences between the WT and cKO (e.g., WT MGE cell population is increased compared to the cKO, but CGE population is decreased in the cKO). **f** Single R analysis results using E16.5 scRNA-seq data from Fig. 2 projected to the adult scRNA-seq public data ^93^ **g** Inferring differentiation trajectories by RNA velocity analysis. The overlaid arrows indicate inferred developmental trajectory. **h** Transcriptional network analysis using scRNA-seq data of the WT and cKO (*Gad2Cre*).

Two single-cell datasets were aligned to compare cell clusters between WT and cKO (Fig. 3c), followed by composition analysis (Fig. 3e). The mutant samples showed a lower proportion of CGE- and a higher proportion of MGE-clusters, as well as subtype alterations in the LGE clusters, with a decrease in D2 MSN, an increase in D1 MSN Tshz, and no detectable changes within the D1 MSN Pdyn cluster (Fig. 3e). Single R analysis that projected our E16.5 scRNA-seq data onto published adult data, demonstrates changes in ratios of different IN subtypes in cKOs (e.g., increase in SST chodl cells) (Fig. 3f) ^61^.

To gain further insights into the cell differentiation, we performed RNA velocity analysis which can predict the direction and speed of cellular state transitions from scRNA-seq data ^62^. Compared to a clear and well-defined differentiation pseudo time flow mapped in the WT MGE cluster (Fig. 3g), a broken MGE trajectory in the cKO indicated an impaired or disrupted differentiation process ^62^. Furthermore, while the WT cells exhibited distinct differentiation patterns between the MGE and CGE, without any connected trajectories, the cKO samples displayed a continued differentiation line without clear boundaries. Together these findings suggest that the loss of *Arx* in the *Gad2*-lineage cells impact their proper differentiation, particularly in MGE- and CGE-derived cINs.

Next, we performed gene regulatory network (GRN) analyses by taking advantage of CellOracle, a machine learning-based approach that combines GRN models (inferred from single cell transcriptomics data plus promoter and TF-binding information) with transcription factor perturbation to simulate the consequences after KO or overexpression of transcription factors ^51^. Using a base GRN built into the CellOracle library and our scRNA-seq data, we generated WT- vs. cKO-specific GRN configurations for MGE cluster cells (Fig. 3h). In WT, major hub genes identified in MGE include *Sp9*, *Mef2c*, *Pbx3*, *Sp8*, and *Maf* (Fig. 3h and Supplementary Fig. 4b). However, in cKO, some of these genes (e.g., *Maf*) were no longer detected as a major hub gene or with altered weight of influence in a network, as displayed in eigenvector centrality plot (Supplementary Fig 4b). These findings indicate perturbed gene regulatory networks in cKO mice.

### Differentially expressed genes (DEGs) in Arx-deficient Gad2-lineage cells

Our data suggest ARX likely plays key roles in multiple steps of IN development: initiation of differentiation by promoting cell cycle exit, subtype specification/maintenance, and initial entry to the cortex during migration. To establish the ARX-related molecular pathways involved in these aspects of IN development, we first conducted machine learning-based single cell transcriptome analysis (scvi-tools package)^63^ and compared transcriptomic changes in the MGE, CGE, and LGE cell clusters between WT and *Arx*-deficient cells (*Gad2*-lineage). The Venn diagrams demonstrate that the MGE and CGE clusters have more up- or down-regulated genes compared to the LGE (MGE, 1008 up/365 down; CGE, 549 up/194 down; LGE, 342 up/172 downregulated genes; |fold change| > 2; p < 0.05) (Fig. 4d and Supplementary Table 1), possibly reflecting the higher expression of *Arx* in the MGE and CGE (Supplementary Fig. 4a).

**Figure 4.**
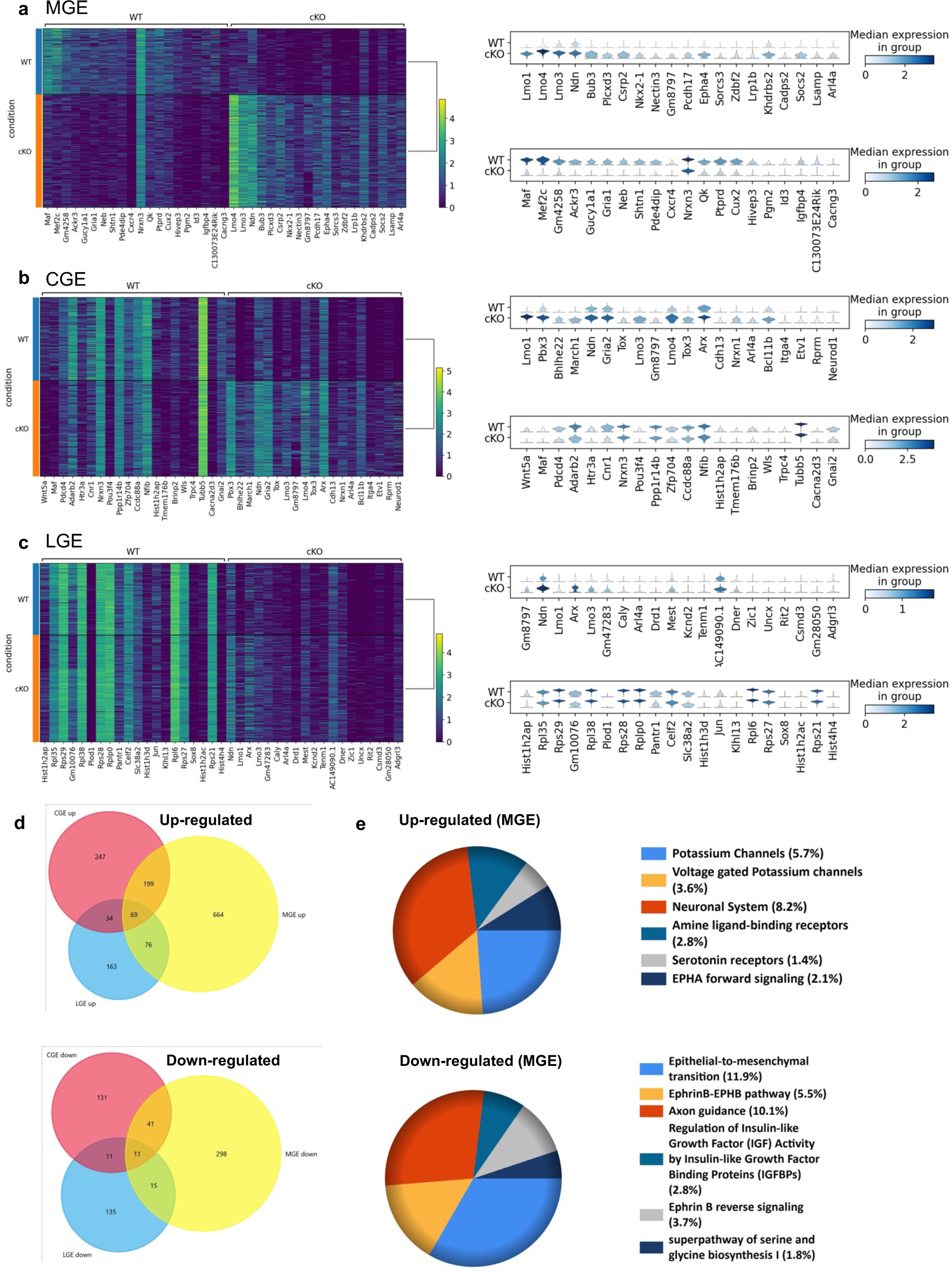
Differentially expressed genes in *Gad2Cre*-*Arx* cKO INs (E16.5?). **a-c** Heatmaps of MGE (a), CGE (b) and LGE (c) clusters displaying differentially expressed genes in cKO (MT), compared to controls (WT). Circled genes are example genes associated with IN progenitor status (red), cell cycle-stemness (green), and ligands/receptors (blue). **d** Venn diagram of up-regulated and downregulated genes in MGE, LGE and CGE INs. CGE, caudal ganglionic eminence; LGE, lateral ganglionic eminence; MGE, medial ganglionic eminence. **e** Gene set enrichment analyses of the upregulated and downregulated genes in MGE INs.

Highlighting the top 20 up- and downregulated genes revealed cell-cycle and cell proliferation associated genes (e.g., *Bub3*, *Csrp2*) ^64,65^, subtype differentiation-associated genes (e.g., *Maf*, *Mef2c*) ^16,17,66^, and cell migration associated genes (e.g., *Cxcr4*, *Ackr3*) ^35,67–70^ in MGE and CGE clusters (Fig. 4a, b). Most of the DEGs in LGE had lower fold changes when compared to MGE and CGE clusters (Fig. 4c). Interestingly, *Lmo1* was upregulated in all three GEs, and a few transcripts implicated in human conditions such as epilepsy, autism spectrum disorder, and Prader-Willi syndrome (e.g., *Ndn*, *Cadps2*, *Lsamp*, *Nrxn*, *Cacng3*) ^71–75^ were also detected in MGE, LGE, and CGE clusters (Fig. 4a, b). Consistent with our DEG analysis, the gene set enrichment analysis (GSEA) identified pathways involved in epithelial-mesenchymal transition (11.9%), axon guidance (10.1%) and EphrinB-EPHB pathway, all of which are frequently associated with cell migration ^22^ (Fig. 4e and Supplementary Table 2). These DEG and GSEA results strongly support that ARX-regulated genes play key roles in cIN generation/differentiation, subtype specification, and migration.

### DEGs play roles in IN differentiation, subtype specification, and migration

To validate the DEGs identified in our scRNA-seq analysis, we performed a series of immunolabeling on *Gad2Cre*-*Arx* cKO and WT brains. Normally, *Nkx2.1* (one of the top 20 upregulated genes) is predominantly expressed in MGE progenitor cells, downregulated as the cells begin to migrate and differentiate, and not detected in the LGE or within the cortex (Fig. 5a-a’’’) ^5,76,77^. Surprisingly, ectopic NKX2.1*+* cells were detected in the cortex, LGE, and CGE of E14.5 cKO brains (Fig. 5b-b’’’ and Supplementary Fig. 5a), and its expression persisted in *Arx*-deficient cINs (tdTomato+) at P0 (Fig. 5d-d’’, f-f’’’), while no WT cINs (tdTomato+) expressed NKX2.1 (Fig. 5c-c’’, e-e’’’). Given the role of NKX2.1 in SST- and PV-expressing IN differentiation ^78^ and its downregulation in postmitotic cells is necessary for IN migration to the cortex ^15^, these data support a failure to down-regulate *Nkx2.1* as the explanation, at least in part, for the observed cIN differentiation and migration phenotypes.

**Figure 5.**
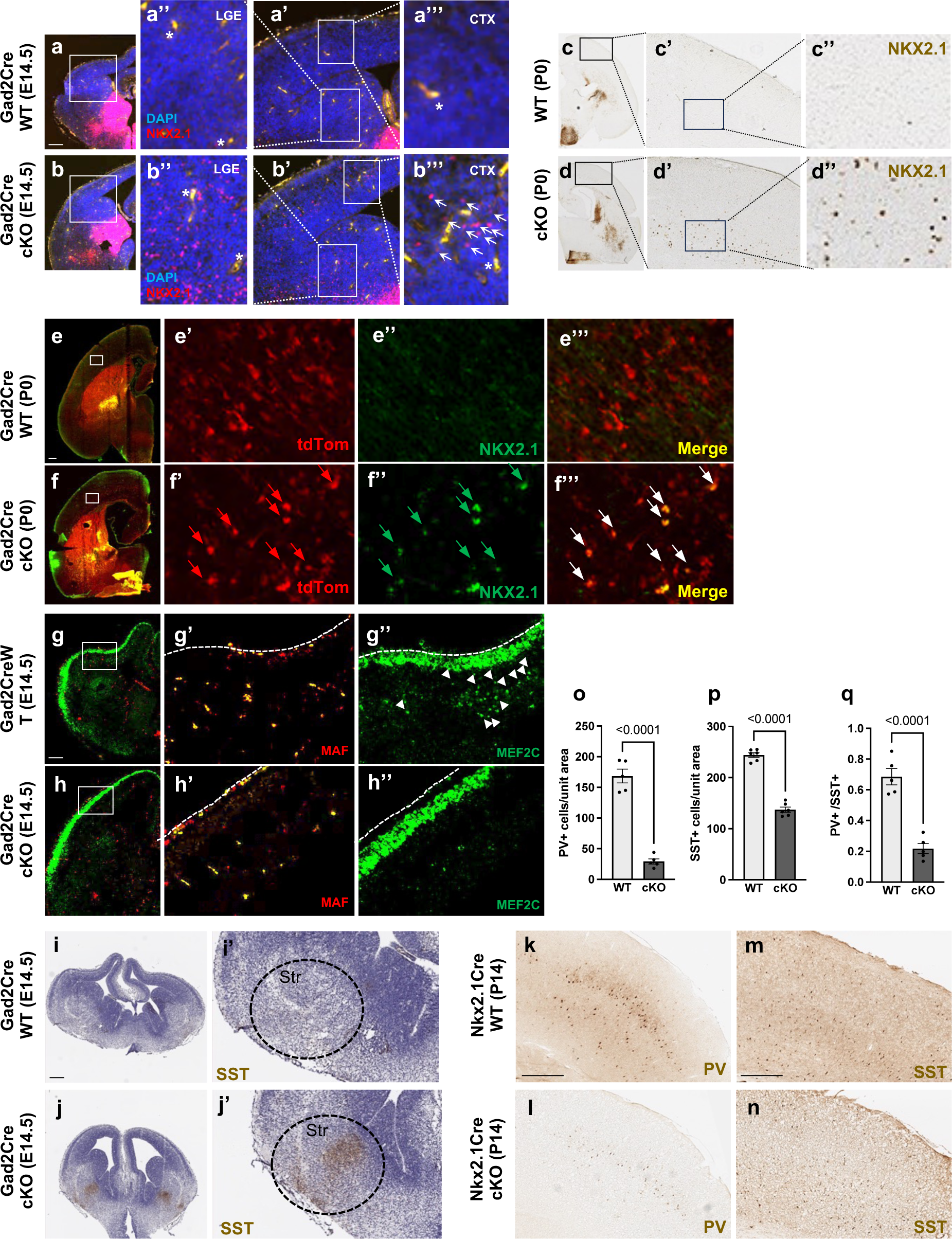
Abnormal IN subtype development in *Arx* cKO. **a-d, e-f** Representative images of NKX2.1 (red) immunolabeling on E14.5 (a-b) or P0 (c-d, e-f) coronal sections of the WT (a, c, e) or *Gad2Cre-Arx* cKO (b, d, f) brains. DAPI (blue) was used for nuclear staining. a’, b’: magnified images of the boxed areas in a, b; a’’, a’’’: magnified images of a’; b’’, b’’’: magnified images of b’; c’, d’: magnified images of c, d; c’’, d’’: magnified images of c’, d’; e’, e’’, e’’’: magnified images of e; f’, f’’, f’’’: magnified images of f; g’, g’’: magnified images of g; h’, h’’: magnified images of h. White arrows in b’’’: NKX2.1+ cells; asterisks: blood cells with autofluorescence; red, green, white arrows in f’, f’’ f’’’, f’’: tdTomato+, NKX2.1+, double+ cells. a-b, e-f: immunofluorescent labeling; c-d: DAB-based immunolabeling. **g-h** Representative images of RNA-scope assay with *Maf* (red) (g, g’, h, h’) or *Mef2c* (green) (g, g’’, h, h’’) probe on E14.5 coronal sections of the WT (g, g’, g’’) or *Gad2Cre-Arx* cKO (h, h’, h’’). **i-j** Representative images of SST immunolabeling (DAB) on WT (i, i’) or *Gad2Cre-Arx* cKO (j, j’) on E14.5 coronal sections. Dotted circles: indicate areas where increased SST expression is increased in cKO.). **k-n** Representative images of PV or SST immunolabeling on WT (k, m) or *Nkx2.1Cre-Arx* cKO (l, n) on E14.5 coronal sections. Dotted circles: indicate areas where increased SST expression is increased in cKO. **o-q** Quantification of PV+ (o) or SST+ (p) IN density and the ratio of PV INs to SST INs (q) from P14 Nkx2.1Cre WT or Arx cKO. n = 6 sections from 2 brains. Scale bars: 200 μm

We next performed immunolabeling for MAF and MEF2C, known to promote PV IN fate and strongly downregulated in the scRNA-seq analysis (Fig. 4a-b) ^16,17^. Immunolabeling confirmed both to be reduced in cINs at E14.5 (Fig. 5g-h, g’-h’, g’’-h’’), predicting a decrease in PV+ INs. As *Gad2Cre-Arx* cKO mice die shortly after birth, prior to PV expression, we were unable to confirm a reduction in PV in these mice. However, *Nkx2.1Cre-Arx* cKO mice, which survive up to P14, show a dramatic decrease in PV+ cINs (Fig. 5k, l, o), in agreement with our prediction (Fig. 4a and Fig. 5g-g’’, h-h’’). Interestingly, SST+ cells, which differentiate earlier than PV+ cells ^79^, were dramatically increased in the cKO striatal region at E14.5 (Fig. 5k-k’, l-l’). In agreement with this, early markers for SST INs were also increased (Supplementary Fig. 5), however by P14 the reduction of SST+ cINs was noted though relatively spared when compared to PV+ cINs (Fig. 5m-n, p-q).

Additional DEGs are known to participate in cell migration (Fig. 4). *Cxcr4* and *Ackr3*, encoding receptors for *Cxcl12* and normally expressed in migrating INs ^35,67–70,80^, were among the top 20 downregulated genes in MGE clusters (Fig. 4). CXCR4 immunolabeling confirmed that its cortical expression in the MZ (arrows in Fig. 6a) and IZ/SVZ (arrowheads in Fig. 6a) were either reduced (MZ) or lost (IZ/SVZ) in the cKO (*Gad2Cre*) (Fig. 6b). Strikingly, its normal expression in the striatum (white arrows in Fig. 6a) was not detected in the cKO brains (Fig. 6b). Given that CXCR4 is a required receptor for IN migration to the cortical MZ ^35,68^, these data provide, at least in part, a molecular mechanism for the deficit of cINs in the cKO MZ stream (Fig. 1b, b’, b’’). In contrast, the IZ/SVZ migration stream was relatively spared, despite the absence of CXCR4 (Fig. 1b’), suggesting the likelihood of redundant or compensatory mechanisms for the IZ/SVZ migration stream.

**Figure 6.**
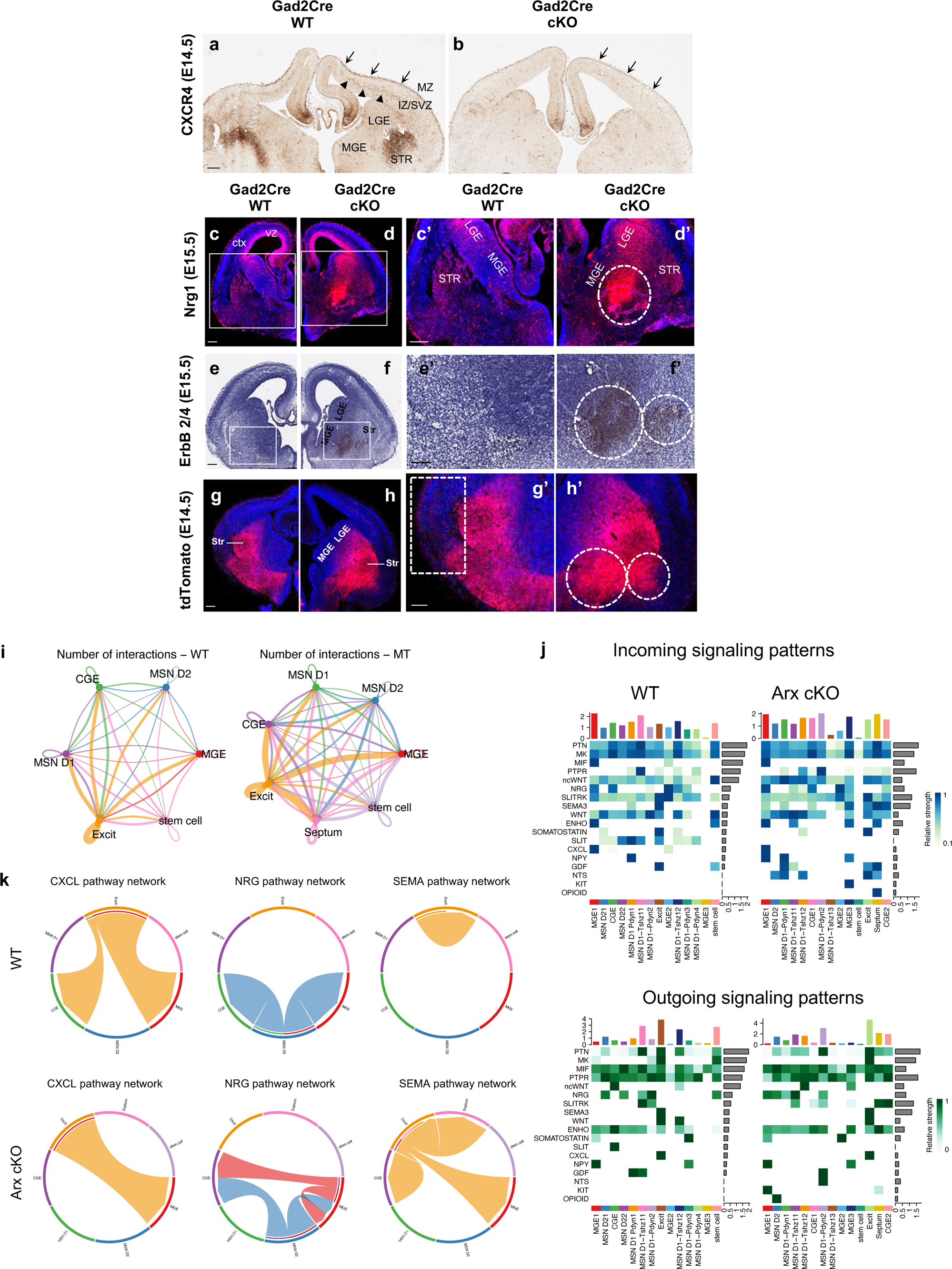
IN migration-associated ligands and receptors are mis-regulated in *Arx* cKO mice. **a-b** Representative images of CXCR4 immunolabeling (DAB) on coronal sections of WT(a) or *Gad2Cre-Arx* cKO (b) brain. Arrows and arrowheads: CXC4+ cells in MZ and in IZ/SVZ, respectively. **c-d** Representative images of *Nrg1* RNA in situ hybridization (RNA scope) on coronal sections of WT(c) or *Gad2Cre-Arx* cKO (d) brain. c’, d’: magnified images of c, d. **e-f** Representative images of ErbB2/4 immunolabeling (DAB) on coronal sections of WT(e) or *Gad2Cre-Arx* cKO (f) brain. e’-f’: magnified images of e-f. **g-h** Representative images of tdTomato immunolabeling on coronal sections of WT (g) or *Gad2Cre-Arx* cKO (h) brain. These sections are the same sections used in Fig.1b-c. **i** Circle plots demonstrating the numbers and strengths of interactions between each cell type identified in single cell transcriptomics of WT and *Gad2Cre-Arx* cKO (MT). Line represents connection between two cell types; line thickness represents interaction strength; loop represents the interaction within the same cell type. **j** Heatmaps displaying representative signals contributing to outgoing or incoming signaling of each cell group identified in scRNA-seq of WT and cKO. **k** Chord diagram depicting all the significant ligand-receptor pairs (cell-cell interactions) between different cell types in WT and cKO.IZ/SVZ: intermediate zone/subventricular zone; LGE: lateral ganglionic eminence; MGE: medial ganglionic eminence; MZ: marginal zone; STR: striatum. Scale bars: 200 μm

Neuregulin1 (*Nrg1*) and its receptor *ErbB4,* both upregulated in the MGE and CGE by scRNA-seq analyses and immunolabeling, are also important for cIN navigation (Fig. 6c-f and Supplementary Fig. 6). Normally, *ErbB4*-expressing INs follow a corridor through LGE that is NRG1+ and lacks SEMA3A/3F, to enter into the cortex ^32,33^. However, ectopic upregulation of *Nrg1* and *ErbB4* in the cKO MGE and CGE raises the possibility that INs are misguided (attracted) to the MGE and CGE. Indeed, tdTomato+ *Gad2*-lineage INs were found to accumulate in the adjacent areas of the striatum (Fig. 6g-h, g’-h’), where ectopic upregulation of *Nrg1* and *ErbB4* were detected (see dotted circles in Fig. 6d’, f’; compare them to those in Fig. 6h’). Furthermore, semaphorin3A (SEMA3A) and its receptor neuropilin1 (NRP1), which normally functions as repellent signaling from the striatum, were upregulated in cKO and likely repel INs from striatum toward ectopic *Nrg1*- and *ErbB4*-expressing MGE areas (Supplementary Fig. 6). Together, these data provide a model where the loss of *Cxcr4* and upregulation of *Nrg1*-*ErbB4* and *Sema3a*-*Nrp1* results in the failure of INs to enter the cortex.

### IN migration defects result from disrupted cell-cell communication networks

To further analyze the significance of changes in ligand-receptor expression, we took advantage of CellChat, a tool used for quantitative inference and analysis of cell-cell communication networks from scRNA-seq data ^50,81^. First, we built a global cell-cell communication networks among the cell types in WT and cKO (*Gad2Cre*) respectively, by counting the number and the strength of interactions and plotting them onto circle-plots according to ligand-receptor pairs (Fig. 6i; see legends for details). Based on these analyses, the number of cell-cell interactions is predicted to be higher in cKO brains than in WT brains (Fig. 6i), suggesting disrupted regulation and chaotic interactions among cKO INs. The strength of the interactions was also higher in cKO brains, probably due to an increased expression of ligands, receptors, or both (as shown in Fig. 6c-f and Supplementary Fig. 6).

CellChat analysis also enables one to predict key incoming and outgoing signals for specific cell types by quantitatively measuring ligand-receptor networks with pattern recognition approaches. As shown in the heatmap of multiple signal strengths (Fig. 6j), each cell type could express specific sets of ligands with distinct strengths, being a signaling sender (see outgoing signaling patterns), and they could also be a signaling receiver if their cell surface receptors are recognized by ligands secreted by other cells or themselves (see incoming signaling patterns). Comparison of both outgoing and incoming signaling patterns highlights the changes in ligand-receptor interaction landscape in each cell type upon *Arx* deletion. The changes in CXCL, NRG, and SEMA signaling are also demonstrated in Chord diagrams showing all significant interactions between different cell types (Fig. 6k). Taken together, the CellChat analyses provide another layer of evidence on disrupted cell-cell communication that results in IN migration defects in cKO brains.

### Regulation of genes involved in IN differentiation and migration by ARX

To determine which genes are directly regulated by ARX, we performed ChIP-seq analysis using E14.5 mouse forebrains (Fig. 7 and Supplementary Table 3). Binding site analyses show ARX binding sites are located near the transcription start site (TSS) and enriched in promoters and exons in the genome (Fig. 7a, b). We also identified enriched consensus binding motifs for ARX (Fig 7c). 607 of the 6590 genes identified by ChIP-seq were also among the 1484 DEGs (Fig. 7d).

**Figure 7.**
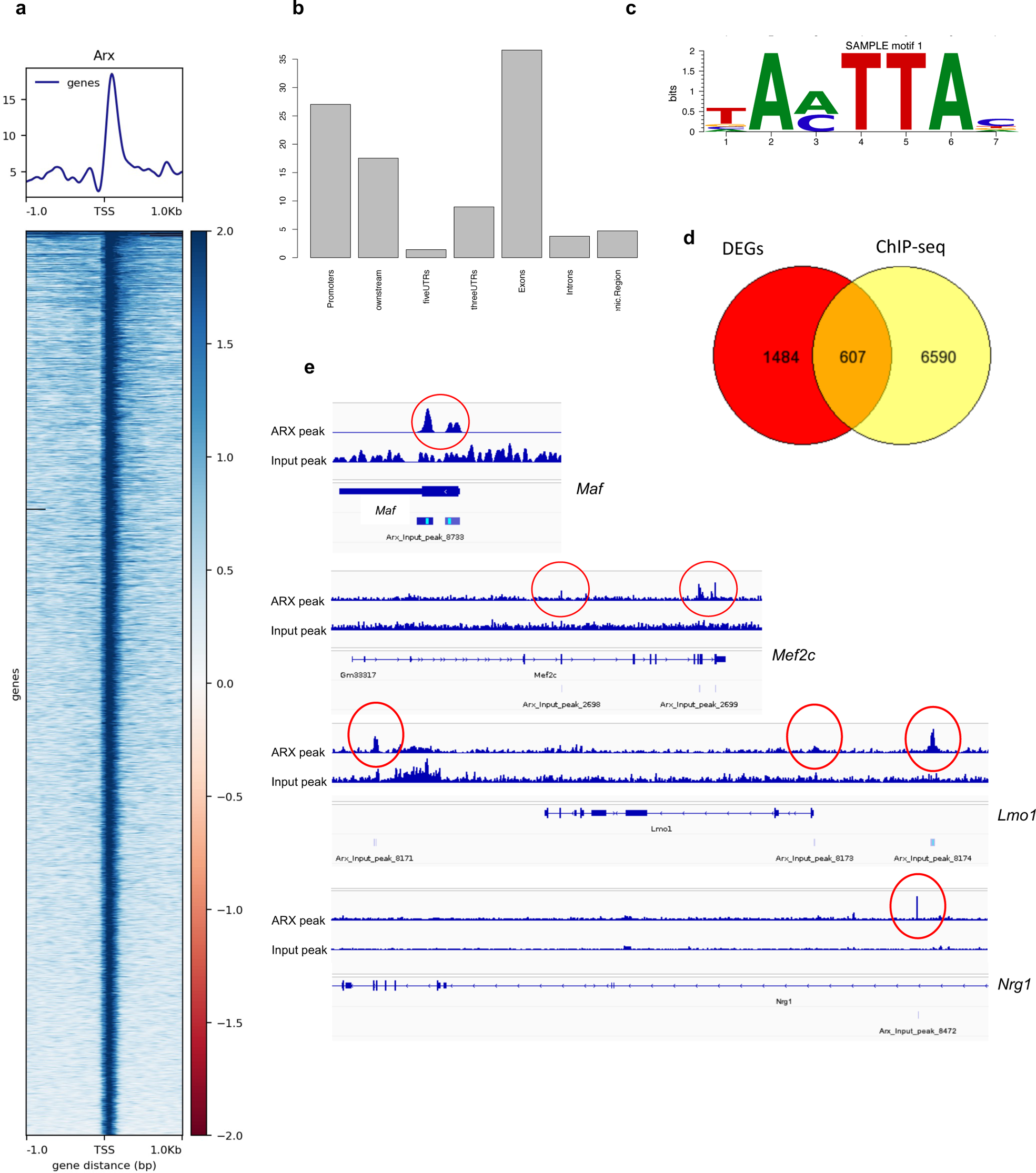
ChIP-seq analysis with anti-ARX antibody. **a** Heatmap plot of ARX binding sites near transcription start site (TSS). **b** ARX binding regions of the mouse genome. **c** ARX binding site sequences in ChIP’ed DNA. **d** Comparison of differentially expressed genes (DEGs) identified in scRNA-seq vs. the genes identified in ARX ChIP-seq. **e** Examples of gene loci (*Maf*, *Mef2c*, *Lmo1* and *Nrg1*) identified as ARX binding sites in ARX ChIP-seq analysis. Red circles indicate examples of ARX specific peaks in each gene loci.

The ChIP-seq data confirmed key DEG loci involved in IN development. For example, *Bub3* and *Csrp2* loci (Supplementary Fig. 7), upregulated genes in the MGE clusters, are known to function in the cell cycle and in the maintenance of stemness ^64,65^. *Maf* and *Mef2c*, downregulated genes in the MGE and CGE clusters direct PV IN subtype specification at the expense of SST INs ^16,17,66^ (Fig. 7e). *Lmo1*, upregulated in the MGE, CGE, and LGE clusters along with *Nrg1* and *ErbB4*, all participate in IN migration ^27,32,33^ (Fig. 7e and Supplementary Fig. 7; also see Fig. 9d, g). Of note, we did not observe an interaction between ARX and the *Cxcr4* locus, however, a previous report identified LMO1 directly interacting with the *Cxcr4* locus (Supplementary Fig. 9c) ^82^, suggesting ARX indirectly regulates *Cxcr4* expression by directly repressing *Lmo1* (also see Fig. 9). In summary, these data are consistent with ARX playing a role in multiple aspects of IN development by directly regulating transcription of a subset of genes involved in the cell cycle, differentiation, subtype specification, and migration, while indirectly regulating other genes.

### Arx^(GCG)7/Y^ mice show cIN density and distribution defects

Studying IN differentiation in *Arx* cKO models is limited as most of the cKO mice we have investigated do not survive until INs fully mature (*Gad2Cre-Arx* cKO die at birth; *Nkx2.1Cre-Arx* cKO die before weaning age; *SstCre-Arx* cKO lives up to 1-2 months). To address this issue and to use a germline mouse model reflecting human patient mutations, we studied *Arx^(GCG)7/Y^* mice ^83^. These mice have an insertional mutation (additional seven alanines) in the first of four poly alanine (PA1, 2, 3, 4) tracks and survive past weaning ^83^. Alanine expansions in PA1 or PA2 represent the most common mutations in patients and have been linked to epilepsy and intellectual disabilities in humans and mice ^84^.

We first examined E14.5 brains for cIN density and distribution using α-DLX antibody ^85^. The density of DLX2+ INs was comparable both in the *Arx^(GCG)7/Y^* and WT (Fig. 8a and Supplementary Fig. 8). The density of DLX2+ INs in the MZ migration stream is lower in the *Arx^(GCG)7/Y^* mice, although not as dramatically reduced as in the cKO models we studied (Fig. 8a, b and Fig. 1a-f). In addition, DLX2+ INs in the MZ and IZ/SVZ migration stream of the mutant cortex appear dispersed, unlike the highly organized streams in WT mice (Fig. 8a).

**Figure 8.**
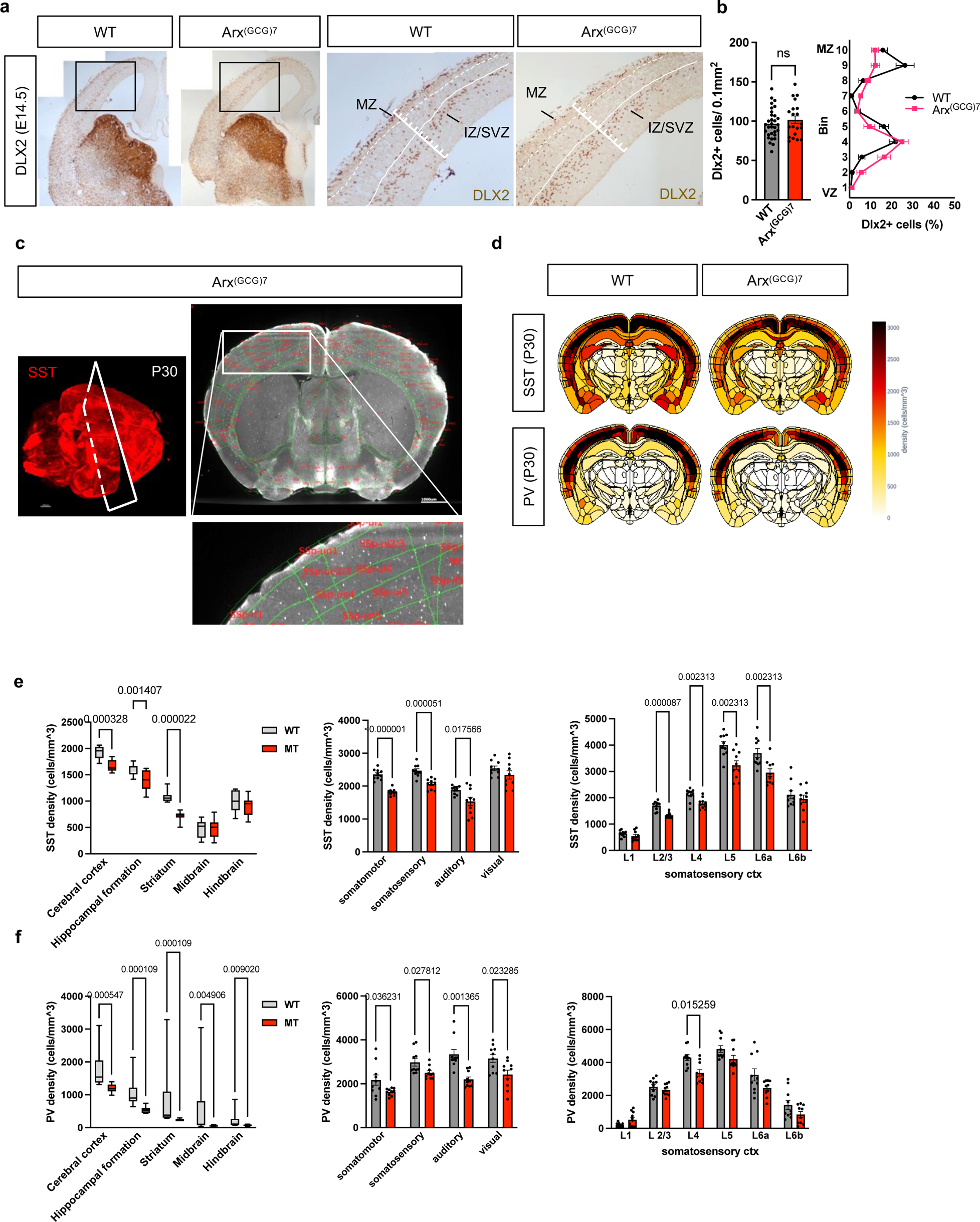
Loss of SST and PV cortical interneurons in Arx^(GCG)7^ mice. **a** Representative images of DLX2 immunolabeling on coronoal sections of the embryonic (E14.5) WT and Arx^(GCG)7^ forebrains. **b** Left: Quantification of the percentage of DLX2+ cells in each bin over the total number of DLX2+ cells in all 10 bins. Right: Quantification of tdTomato+ cell density in WT and Arx^(GCG)7^. **c** Representative analysis of 3D immunolabeling of SST on adult brains (P30) of Arx^(GCG)7^. Left top: a dorsal view of the whole brain with a slice at the level of anterior hippocampus. Left middle: coronal view of the boxed area in the top. Left bottom: magnified image of the boxed area in the middle. Right top: An example coronal section (SST immunolabel) overlayed with reference brain atlas for quantitative analysis. Right bottom: Magnified image of the boxed area in top (red labels and green lines indicate each area annotated in the Alan Brain Atlas). **d** Heat maps on the coronal section illustrations showing the densities of the SST or PV immunolabeled cells of the WT and Arx^(GCG)7^ brains. **e-f** Quantification of SST IN (e) or PA IN (f) cell density (cells/mm^3) in each indicated brain regions (left), each cortical area (middle), or each indicated layer of the somatosensory cortex (right) of the WT and Arx^(GCG)7^(labeled as MT). Scale bars: 200 μm

Leveraging whole brain tissue clearing technique combined with immunofluorescence, we next imaged the distribution of SST and PV neurons in the whole brain at P30 (Fig. 8c) ^23^. The resulting 3D image analyses revealed that the density of SST INs in *Arx^(GCG)7/Y^* mice was reduced in the cerebral cortex, hippocampus, and striatum, while no significant differences were observed in the midbrain and hindbrain (Fig. 8e). Among the neocortical areas, the differences of the SST IN in the motor- and somatosensory cortices were most severe, while differences in the auditory and visual cortices showed no statistical difference (Fig. 8d, e). Layer-specific distributions of SST INs in the somatosensory cortex identified a significant loss of SST INs Layer 2-6a, which is slightly different from the *Arx* cKOs showing preferential loss in upper layers (Fig. 8e). The density of PV INs showed a milder reduction compared to that of SST INs, reaching statistical significance only in Layer 4 (Fig. 8f). VIP INs did not show significant reductions in the brain (Supplementary Fig. 9a). Together these results indicate that *Arx^(GCG)7/Y^* mice show milder cIN defects relative to *Arx* cKO mice with several unique features.

### Lmo1 and Cxcr4 contributes to cIN migration defects in Arx mutant mice

To further understand the molecular basis of cIN phenotypes in *Arx^(GCG)7^* mice, we performed an RNA microarray analysis. *Arx^(GCG)7^* mice were crossed with *Dlx5/6CreGFP* transgenic line to label DLX5/6+ INs with GFP. GFP+ cells were isolated from E14.5 *Arx^(GCG)7/Y^*;*Dlx5/6CreGFP* and *Arx^+/Y^*;*Dlx5/6CreGFP* mice to identify DEGs between *Arx^(GCG)7/Y^*and WT mice. We compared the list of up- and downregulated genes in *Arx^(GCG)7/Y^* GE cells with those in *Arx* constitutive KO GE cells (E14.5) (Fig. 9b and Supplementary Table 4) ^21^. As anticipated, *Arx^(GCG)7/Y^* GE cells had fewer mis-regulated genes (with fold change >1.5 and adj-P value <0.05), compared to *Arx*-deficient GE (*Arx* KO). Among the 45 total upregulated genes in *Arx^(GCG)7/Y^* GE, 15 of them (including *Lmo1* and *Lmo4*) overlapped with those in *Arx* KO cells (total 1329 genes were upregulated). Similarly, among 22 downregulated genes in the *Arx^(GCG)7/Y^* GE, only *Cxcr4* and *Lhx8* were also downregulated in *Arx* KO GE cells (total 845 genes downregulated) (Fig. 9b). We confirmed the downregulation of *Cxcr4* (an essential chemokine receptor gene for cIN migration) (Fig. 9c).

**Figure 9.**
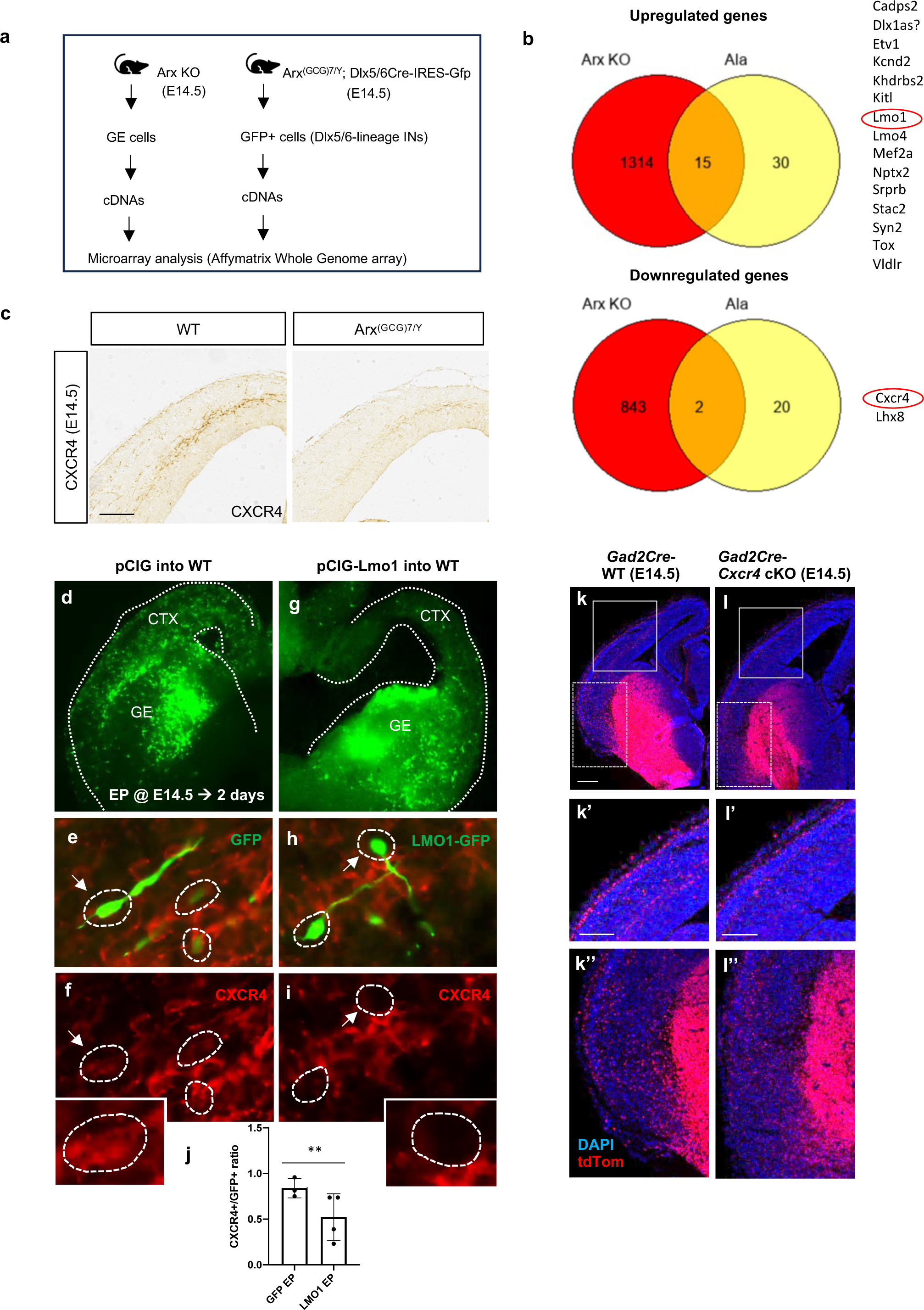
Loss of CXCR4, due to upregulated *Lmo1* expression, contributes to IN migration defects in Arx^(GCG)7^. **a** Summary schematic depicting microarray analysis in Arx^(GCG)7^; Dlx5/6CreGFP and Arx KO mice (E16.5?). **b** Venn diagram of upregulated and downregulated genes identified from RNA microarray analysis of the *Arx* KO vs. Arx^(GCG)7^ GE tissues, compared to each WT littermate set. **c** Representative images of CXCR4 immunolabeling on coronal sections of the embryonic (E14.5) WT and Arx^(GCG)7^ forebrains. **d, g** Forced expression of pCIG (d) or pCIG-Lmo1 (g) into the GE of WT brain slice cultures via electroporation at E14.5 (analyzed 2 days later). **e**-**f**, **h**-**i** Migrating INs electroporated with pCIG (green cells marked with dotted circles in e) expresses Cxcr4 (red in f), but those with pCIG-Lmo1 (green cells marked with dotted circles in h) lost Cxcr4 expression (note no signal in dotted cells in i). **j** Quantification of the ratio of CXCR4+ cells among GFP+ cells in pCIG or pCIG-Lmo1 electroporated slice culture. Error bars: mean ± sem. **k-l** Representative images of tdTomato immunolabeling on coronal sections of the WT (k) or *Cxcr4* cKO mediated by *Gad2Cre* (WT and Cxcr4 cKO mice are on Ai14 background). k’ and l’ are magnified images of the cortical areas in k and l, respectively. Scale bars: 200 μm

As *Lmo1* has been consistently detected as a differentially expressed gene in multiple transcriptomic analyses of *Arx*-deficient as well as *Arx^(GCG)7/Y^*mouse brains (Fig. 4a, b and Fig. 9b; ref. ^42,86,87^), and it has been shown to bind to *Cxcr4* loci (Supplementary Fig 9c) ^82^, we explored its function by overexpressing *Lmo1* in the GE of WT mice. IN migration progressed normally in brain slice cultures electroporated with a GFP construct (*pCIG*) into the GE (Fig. 9d). In contrast, most INs electroporated with the *Lmo1* construct (*pCIG-Lmo1*) failed to migrate to the cortex (Fig. 9g). These data provide evidence that the upregulated *Lmo1* in *Arx*-mutant INs, at least in part, responsible for the IN migration/distribution defects observed in *Arx* mutant mice (both cKO and *Arx^(GCG)7/Y^*). Interestingly, when these slice cultures were immunolabeled with an anti-CXCR4 antibody, migrating INs electroporated with *Lmo1* did not express *Cxcr4*, while those electroporated with GFP showed normal *Cxcr4* expression (Fig. 9e, f, h, i, j). These findings are consistent with the significantly reduced *Cxcr4* expression in *Arx* cKO as well as *Arx^(GCG)7/Y^* mice (Fig. 6a, 9c). To further test this, we generated *Gad2Cre*-*Cxcr4* cKO lines crossed with Ai14 to abrogate *Cxcr4* in *Gad2*-lineage INs while simultaneously labeling *Cxcr4*-abrogated cells with tdTomato. Interestingly, *Cxcr4* cKO mice also showed a similar cIN migration phenotype observed in *Arx^(GCG)7/Y^* mice (Fig. 9k, l, k’, l’) ^70^, suggesting that the downregulation of *Cxcr4* by abnormally upregulated *Lmo1* is likely to be responsible for the cIN migration phenotype in *Arx^(GCG)7/Y^* mice. Tissue clearing combined with 3D image analysis revealed no significant changes in PV+ and SST+ IN densities in *Cxcr4* cKO mice (Supplementary Fig. 9b, c). Another receptor, CXCR7 (known as ACKR3) also binds to CXCL12, and a similar result was obtained with *Gad2Cre*-*Cxcr7* cKO lines crossed with Ai14 (Supplementary Fig. 9d).

## Discussion

ARX is one of the key transcription factors for cIN development and its mutations are associated with several neurologic disorders ^88^. Elucidating the role of ARX and its associated transcriptional networks is essential to better understand these conditions. Our findings provide evidence that ARX modulates multiple aspects of cIN development and migration, including the transition from proliferation to differentiation, subtype specification, and entry into the pallium and cortical plate.

EdU injection experiments demonstrate that loss of *Arx* from the GE SVZ lowers cell cycle exit rate, likely interrupting the transition from proliferation to differentiation. This is in accordance with abnormal upregulation of *Csrp2*, known to promote stemness ^65^, and *Bub3*, a cell cycle checkpoint protein often overexpressed in tumor cells ^64^. These two genes are direct transcriptional targets of ARX (Fig. 7) and their upregulations in *Arx*-deficient IN precursors likely keep them in proliferation status, inhibiting differentiation. These findings suggest that ARX normally promotes differentiation of IN precursor cells while also inhibiting their cell cycle progression, as reported with other transcription factors involved in differentiation ^89^. Interestingly, this is different from what we observed in the cortical PN precursor cells, where ARX normally represses cell cycle inhibitor, *Kip2* expression, thus suppressing differentiation ^90^. These distinct functions of ARX in cortical vs. GE precursor cells are also reflected in its distinct expression pattern in these two areas: in dorsal (cortical PN) progenitors, it is mainly expressed in the VZ, whereas in the GE ARX is primarily in the SVZ and differentiated cells ^40^.

Our single cell transcriptomic data combined with immunolabeling analyses demonstrate that ARX also impacts IN subtype specification. *Arx* depletion from IN precursors leads to a significant shift, increasing the proportion of MGE derived INs while decreasing those originating from the CGE. The ectopic NKX2.1+ cells detected in CGE of *Gad2Cre-Arx* cKO mice likely accounts for this transition by specifying CGE located cells to become phenotypically MGE. The alteration in cell types was also demonstrated in the single cell analyses from *Gad2Cre-Arx* cKO brains ^61^ where the abnormal upregulation of *Sst* and downregulation of *Maf* and *Mef2c*, early markers of PV INs that promote PV fate and direct transcriptional targets of ARX, were found. These changes in embryonic brains resulted in the reduced ratio of PV+ to SST+ INs in P14 cKO brains. The increase in SST+ cells over PV+ cells raises the possibility of a fate shift of PV+ cells to SST+ cells, similarly to that observed in *Mafb* and *c-Maf* double cKO mice ^16^. Consistent with these data, the early PV IN markers (e.g., *Mef2c*, *Maf*, *Angp2*, *Crbp1*, *St18*) are decreased in cKO MGE clusters, while early SST IN markers (e.g., *Sst*, *Tspan7*, *Cacna2d3*, *Nxph1*) are increased. Interestingly, these *Arx* cKO cell phenotypes resemble those of the *Sp9* KO mice ^91^. Given that Arx is directly regulated by SP9, there is likely biologic pathways connecting *Arx* and *Sp9* ^91^.

Consistent with *Sst* upregulation in *Arx*-deficient cells (in *Gad2Cre-Arx* cKO mice), *SstCre*-mediated *Arx* cKO mice have an increased density of tdTomato+ INs (SST lineage) compared to WT mice, unlike *Gad2Cre*- or *Nkx2.1Cre*-mediated cKO, where tdTomato+ IN densities are lower than those in the WT mice. We speculate that the loss of ARX promotes SST expression, thus facilitating production/differentiation/survival of SST+ cINs in the cortex and not an increase in the migration of cINs from the GE.

The dramatic loss of *Arx*-deficient cINs, specifically in the MZ stream, provides us with unique opportunities to study the biology of IN migration. For example, *Nkx2.1* is a direct transcriptional target of ARX (Supplemental Fig. 7) and one of the top 20 upregulated genes in MGE cluster. Considering that downregulation of *Nkx2.1* in postmitotic cINs is necessary for them to enter cortex migration streams ^15^, we speculate the observed IN migration defects in *Arx* cKO mice may be attributed, in part, to failure in downregulating *Nkx2.1*. In addition, whether NKX2.1+ cells detected in the LGE (in addition to the cortex) are cells enroute to the cortex from the MGE, or are born in the LGE VZ, is not clear and will require further investigation.

Another direct target of ARX involved in cIN migration is *Lmo1*. It is consistently upregulated in *Arx* cKO, constitutive KO, and *Arx^(GCG)7^* mice. Interestingly, LMO1 directly represses *Cxcr4* expression based on ChIP-seq and our slice culture electroporation data. The loss of CXCR4 in *Arx*-deficient INs is thought to be responsible, at least partially, for the failure of INs to enter into the cortical MZ, combined with the ectopic upregulation of the guidance signal Nrg1/ErbB4 (direct targets of ARX). It is not clear how upregulated Sema3A/Nrp1 contributes to the IN migration phenotype in *Arx* cKO mice, but future studies should assess the roles of Nrg1/ErbB4 as well as of Sema3/Nrp1 in directing cINs out of the GE to the cortex, in the context of ARX function.

Finally, we note that previously published studies reported no significant changes in cIN numbers or distribution in *Arx* poly-alanine expansion mouse models, distinct from our data ^92^. We surmise this is due to differences in methodologies; the use of 3D image analysis of tissue cleared, immunolabeled brains permits the counting of all labeled neurons rather than subsets, thus likely a more sensitive measure.

In summary, we have provided evidence that ARX is a hub transcription factor with key roles in the generation, fate specification, and migration of cINs through the regulation of gene transcription networks during brain development. ARX controls multiple steps of IN differentiation by directly or indirectly regulating other genes such as *Bub3*, *Cspr2*, *Maf*, *Mef2c, Nkx2.1, Lmo1*, *Cxcr4*, *Nrg1*, *ErbB4*, *Sema3*, *Npr1*, and *EphA4*. Together our findings provide novel insights on the impact of the ARX on cIN development, and how those roles are disrupted in disease.

## Methods

### Mice

All procedures and animal care were reviewed, approved, and performed in accordance with the Cedars-Sinai Medical Center Institutional Animal Care and Use Committee (IACUC) guidelines. All mouse strains have been previously published: *Ai14* (Rosa 26-tdTomato Cre-reporter) (Jackson Lab #007914) ^43^, *Gad2-IRES-Cre* (Jackson Lab #028867; referred to as *Gad2Cre* in this study) ^44^, *Nkx2.1-CreER* (Jackson Lab #014552)^44^, *Nkx2.1-Cre* (Jackson Lab #:008661) ^45^, *Sst-IRES-Cre* (Jackson Lab #013044; referred to as *SstCre* in this study) ^44^, *Dlx5/6-Cre-IRES-Gfp* (referred to as *Dlx5/6Cre-Gfp*) in this study ^46^, *Arx-flox*^42^, *Arx*^(GCG)7^(also referred to Arx^PA1–22^)^18^, and *Cxcr4-flox* (Jackson Lab# 008767) ^47^. Mice were maintained on a C57BL/6J background. For timed pregnancies, noon on the day of the vaginal plug was designated as embryonic day 0.5. For tamoxifen induction, 3.75 mg of tamoxifen was administered to pregnant dams by IP injection (150 μL of 25 mg/mL stock plus 150 μL of corn oil) and cesarean section was performed at E19.5-20.5 for P0 harvest.

### Single-cell RNA sequencing, differential expressed genes (DEG) analysis and gene set enrichment analysis (GSEA)

E16.5 *Gad2Cre-Arx* cKO and WT forebrains were harvested and incubated with papain solution (Worthington Biochemical) for 20 min at 37°C. After incubation, the single cells were prepared by gentle trituration. TdTomato+ cells were isolated by cell sorter (BD FACSymphony S6) and loaded onto a 10X genomics single-cell preparation platform following the manufacturer’s instruction (10x Genomics). A minimum of 5,000 cells were recovered per sample (2 wild types and 3 mutants, males). Library preparation was performed using 10X Chromium Single 3’ Reagent Kits Version 2 (10x Genomics). The library length and concentration were quantified using a Bioanalyzer (Agilent) according to the manufacturer’s protocol. Paired-end sequencing was performed on the Illumina NovaSeq. FASTQ files were aligned with mouse genome (mm10) by Cellranger V6.0 (10xgenomics). As a result, the cell barcode, UMI code and expression matrix obtained were used for analyses. Using the single-cell variational inference tools (scvi-tools) package ^48^, normalization, removal of doublet, dimensionality reduction, integration across biological conditions, differential expression (DE), and UMAP visualization were performed. Gene set enrichment analysis was performed by FunRich package ^49^ with DE genes filtered with adjusted p-value less than 0.05 and |FC| > 1.5. Cell-cell communication analysis was performed using the R package CellChat ^50^ with default parameters. We ran the python package CellOracle^51^, to compare gene regulatory network models. To visualize the network, we used Gephi ^52^.

### Microarray analysis and NGS transcriptome data analysis

cDNA was amplified from extracted RNA using the QIAGEN RNAeasy kit after purification of GFP-positive interneurons from the forebrain of E14.5 of the *Arx*^(GCG)7/Y^ and WT littermate controls crossed with *Dlx5/6-Cre-IRES-Gfp*. For each microarray, 1 μg of cDNA was labeled and hybridized to an Affymetrix Mouse Whole Genome 430 2.0 array, with six independent replicates per condition. The Bioconductor software package was used to background-correct and normalize the data and calculate mean fold change (FC) and false discovery rates (FDR) for all pair-wise comparisons.

### Chromatin immunoprecipitation and sequencing (ChIP-seq)

Embryonic forebrains (E14.5) were isolated and fixed in 4% fresh paraformaldehyde in PBS for 16 hrs at 4°C. Fixation was quenched with 125 mM Glycine at room temperature for 5 min and washed with cold PBS twice. Chromatin immunoprecipitation was performed using Simplechip® Plus Sonication Chromatin IP Kit (Cell Signaling Technology) with minor modifications as follows. First the forebrain tissue was soaked with citrate antigen retrieval buffer and heated for 15min by double boiling. After antigen retrieval, the tissue was washed with water several times and neutralized with the ChIP buffer. Chromatin was fragmented into 100–300 bp fragments using the Bioruptor UCD-200 (Diagenode; high power) for 2 hr (cycled at 30 sec ON/30 sec OFF). For immunoprecipitation of ARX-bound chromatin, 2 μg of anti-ARX antibody (R&D) was incubated with cleared chromatin lysate. Input and ChIPed DNA libraries were prepared using an Illumina Next Seq (single-end reads of 75 bp). FASTQ sequences were aligned to the mouse mm10 genome sequence using HISAT2 and converted to SAM and then BAM files. ChIP-seq peaks were called using MACS2, input DNA without ChIP as reference using default settings. The HISAT2-aligned peaks and MACS-determined peak positions were visualized using IGV Genome Browser.

### Immunolabeling

Embryonic brains were harvested and fixed in 4% paraformaldehyde (PFA) overnight at 4°C. Postnatal brains (P0 and P30) were harvested after trans-cardiac perfusion (first with PBS containing 60 mM of heparin and then with 4% PFA) and were post fixed in 4% PFA overnight at 4°C. Harvested brains were transferred to 30% sucrose for cryoprotection and stored at 4°C overnight or until ready to use. Brains were embedded in OCT and cryo-sectioned coronally at 14 μm (on a Leica CM1950 cryostat). Immunofluorescence labeling was performed as previously described ^53^. All primary and secondary antibodies were diluted in a non-animal blocking solution (Vector lab). Slides were incubated with primary antibodies overnight, while biotinylated or florescence-conjugated secondary antibodies were incubated for 1 hr at room temperature (RT). Primary antibodies used included rabbit anti-RFP (1: 500, Rockland), rabbit anti-CXCR4 (1: 200, Abcam), rat anti-somatostatin (1: 50, Scbt), anti-Erbb2/4 (1:1000, Abcam), and anti-Dlx2 (1:200, gift from Dr. David D. Eisenstat at University Melbourne). Secondary antibodies conjugated with Alexa-488, -594, or -647 fluorophore were all used at 1:250 (ThermoFisher). Sections were mounted with Prolong Glass Mounting Media (Invitrogen, P36984) containing DAPI (Vector labs). For DAB staining, the ABC kit (Vector lab) and detection system were used according to the manufacturer’s recommendation.

### Cell cycle analyses

Ethynyl deoxyuridine (EdU; Invitrogen) was administered intraperitoneally (50 mg/kg) to pregnant mice (E13.5 for 1.5 hour to establish proliferation rates, 24 hours to determine cell cycle exit, or for 48 hours to follow GE to cortex migration). Click-iT EdU cell proliferation kit (Invitrogen) was used to label EdU+ cells. Cells in active cell cycle were labeled with a Ki67 (D3B5) antibody (1:200 dilution, Cell Signaling Technology).

### RNA scope

Cryosections (E14.5), 16 μm thick, were prepared as previously described^53^. Sections were air dried at 60°C for 1 hr, followed by 4% PFA post-fixation at 4°C for 15 min. Then, target retrieval was performed by double boiling the sections for 7 min and treating with Protease III for 10 min. *In situ* hybridization was performed following the manufacturer’s (ACDbio) protocol for the multiplex fluorescent kit (version 2). RNAscope probes were Mm-Maf (Cat. 412958) and Mm-*Mef2c* (Cat. 421011). Slides were mounted with Prolong glass mounting media (Invitrogen, P36984) containing DAPI.

### Tissue clearing, immunolabeling, and cell counting

P0 and P30 mice were perfused and fixed as described above. Tissue clearing was performed using the SHIELD protocol by LifeCanvas Technologies or iDISCO ^54,55^. For the first method, samples were cleared for 7 days with Clear+ delipidation buffer and then immunolabeled using SmartLabel. Each sample was labeled with 5μg mouse anti-NeuN (RRID:AB_2572267), 4μg rabbit anti-parvalbumin (Swant PV27; 1:400), and rat anti-somatostatin (Millipore) followed by fluorescent secondary antibodies. Samples were incubated in EasyIndex for a refractive index of 1.52 and imaged at 3.6X using SmartSPIM microscopy. Images were tile-corrected, de-striped, and registered to the Allen Brain Atlas (https://portal.brain-mapping.org). The NeuN channel was registered to the 8-20 atlas-aligned reference samples using successive rigid, affine, and b-spline warping (SimpleElastix: https://simpleelastix.github.io). Average alignment to the atlas was generated across all intermediate reference sample alignments to serve as the final atlas alignment value per sample. Fluorescent measurements from the acquired images were projected onto the Allen Brain Atlas to quantify the total fluorescence intensity per region defined by the Allen Brain Atlas. These values were then divided by the volume of the corresponding regional volume to calculate the intensity per voxel measurements. Data was plotted using Prism (version 10).

### Slice culture and electroporation

CD-1 mice were time-mated (plug=0.5 days) and embryos collected on E14.5. The brains were harvested and embedded in 3% low melting point agarose (Fisher Scientific cat.# BP165-25) and sectioned at 300 μm (Leica VT 1000s) ^56,57^. pCIG and pCIG-Lmo1 plasmids were micropipetted into the medial ganglionic eminence of each slice followed by electroporation (Nepa Gene CUY21 electroporator; 60V, pulse interval: 5 ms, pulse duration: 200 ms, number of pulses: 5). Slices were cultured on laminin and poly–L–lysine coated transwell inserts (BD Falcon cat. # 353102) for 3 days at 37°C and 5% CO2 ^58^ after which slices were fixed with 4% paraformaldehyde overnight at 4°C.

### Explant culture and cell migration analysis

Mouse embryonic medial ganglionic eminence (MGE) from E14.5 was dissected in ice-cold HBSS (Sigma) and used for explant culture as described previously ^56^. Briefly, one hundred micrometer square explants were cut out of MGE and cultured in a 3D extracellular matrix gel on a glass bottom 6 well plate (MatTek Corporation). The explants were covered with 50% Matrigel (Corning) and 50% collagen (2 mg/ml; BD Biosciences), and placed at 37°C, 5% CO2 for 20 min to set. Tissue was incubated in DFS media (F12:DMEM w/10% fetal bovine (FBS) serum, 40 μM L-glutamine, and 47 mM glucose, P/S) for 1 h, then switched to DM media (DMEM w/N2, 36 mM glucose, and P/S). The explants were imaged on Zeiss Observer Z1 inverted microscope.

### Image analyses and quantifications

For cell density determination, coronal sections were taken from the levels covering from the septum to CGE. The entire cortical area spanning from the dorso-medial apex to the pallial-subpallial boundary were taken as the ROI. The total numbers of tdTomato+, PV+, SST+, and DLX+ cells in this ROI were counted manually in Image J using StarDist 2D Plug-in, and divided by the area of the ROI (in μm^2^). cIN distribution was established using the entire pallium divided into 10 bins from the ventricular surface (bin 1) to the pial surfaces (bin 10), and a single counting field (200 μm wide for E14.5 and 400 μm wide for P0) per section was used. For quantification of the EdU injection experiments, cortical-GE boundary area was taken as ROI at multiple levels of coronal sections covering from the septum to the CGE. Using StarDist 2D Plug-in, each DAPI+ cell was set per ROI and the number of EdU+ or Ki 67+ cells were counted.

### Quantification and statistical analyses

All statistical analyses were performed using Graphpad Prism (version 10). The analysis of data normality was determined by Anderson–Darling, D’Agostino-Pearson, Shapiro–Wilk, and Kolmogorov–Smirnov tests. To determine statistically significant differences between two data sets, a two tailed t-test (with Welch’s correction) was used if the data passed a normality test; a non-parametric test (Mann Whitney test) was used if the data failed normality test. The details of all statistical tests including the p-values and sample numbers (n) are listed in the figure legends or directly in the graphs. All samples were randomly selected, and image quantifications were carried out blindly. P values of ≤ 0.05 were considered statistically significant. All bar graphs are plotted as mean ± sem and all scattered plots are marked with individual data points and median with inter quartile range.

## Acknowledgements

We are grateful to Dr. David D. Eisenstat (University Melbourne) for providing an anti-DLX2 antibody. We thank Cedars-Sinai Biobank and Research Pathology Resource Core for imaging service and assistance. This work was supported by NIH NINDS grants (R01 NS100007 and R01 NS113516).

## Author Contributions

Y.L., J.A.G, and G.C. conceived and designed the study. Y.L., S.K.A, A.K.M, and G.C collected data. Y.L. and G.C analyzed the data, prepared the figures, and wrote the manuscript together with J.A.G. S.K.A. and C.A.W. contributed to single cell RNA-seq data collection and analysis. C.C. and K.A.R provided technical service and participated in data collection and analysis. All authors reviewed the manuscript.

