## Supplemental Figures 1 to 9 for "ARX regulates cortical interneuron differentiation and migration"

### Supplementary Figure 1

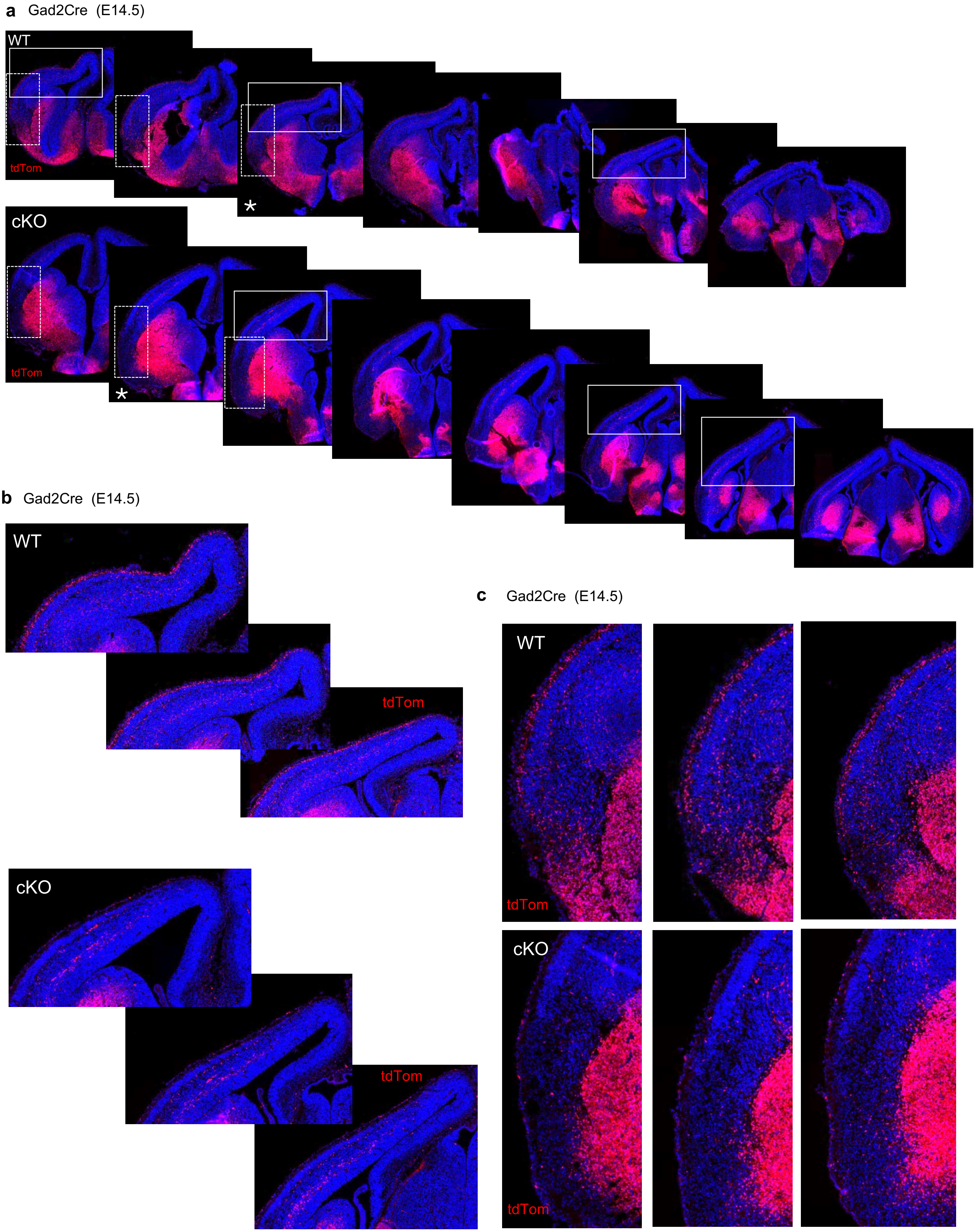

**Supplementary Fig 1**. **a** A representative series of coronal sections from *Gad2Cre-Arx* cKO or WT brains (E14.5) immunolabeled with tdTomato (red) and nuclear stained with DAPI (blue). Asterisks indicate the same sections shown in Fig. 1b, c. **b** Magnified images of solid boxed areas in a. **c** Magnified images of dotted boxed areas in a.

### Supplementary Figure 2

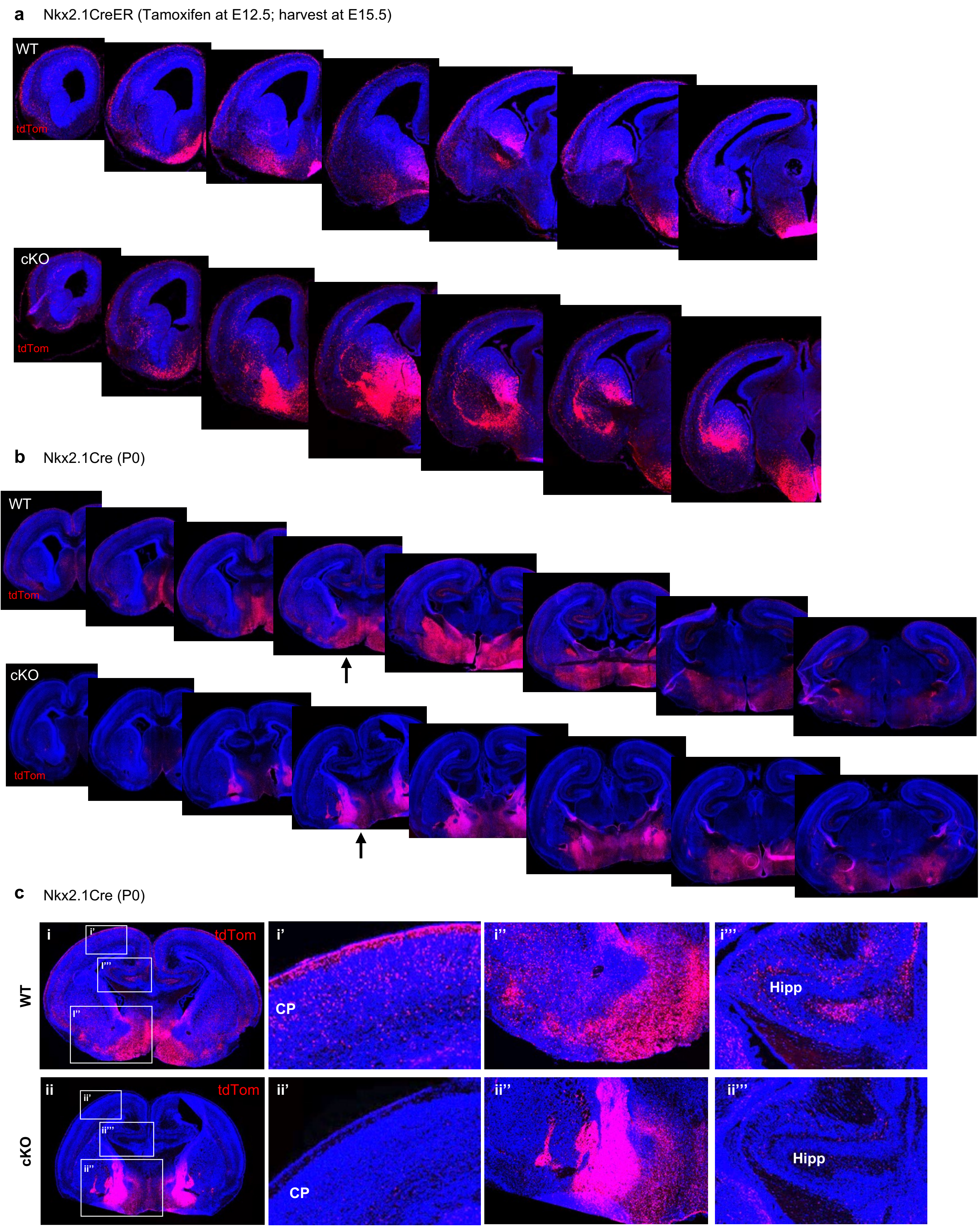

**Supplementary Fig 2**. **a** A representative series of coronal sections from *Nkx2.1CreER-Arx* cKO or WT brains (E15.5) immunolabeled with tdTomato and nuclear stained with DAPI (blue). **b** A representative series of coronal sections from *Nkx2.1Cre-Arx* cKO or WT brains (P0) immunolabeled with tdTomato. **c** Representative images of Nkx2.1Cre-Arx cKO or WT (P0) brain sections marked with arrows in b. I', ii', and iii' are magnified images of boxed areas in i. CP, cortical plate; Hipp, hippocampus.

### Supplementary Figure 3

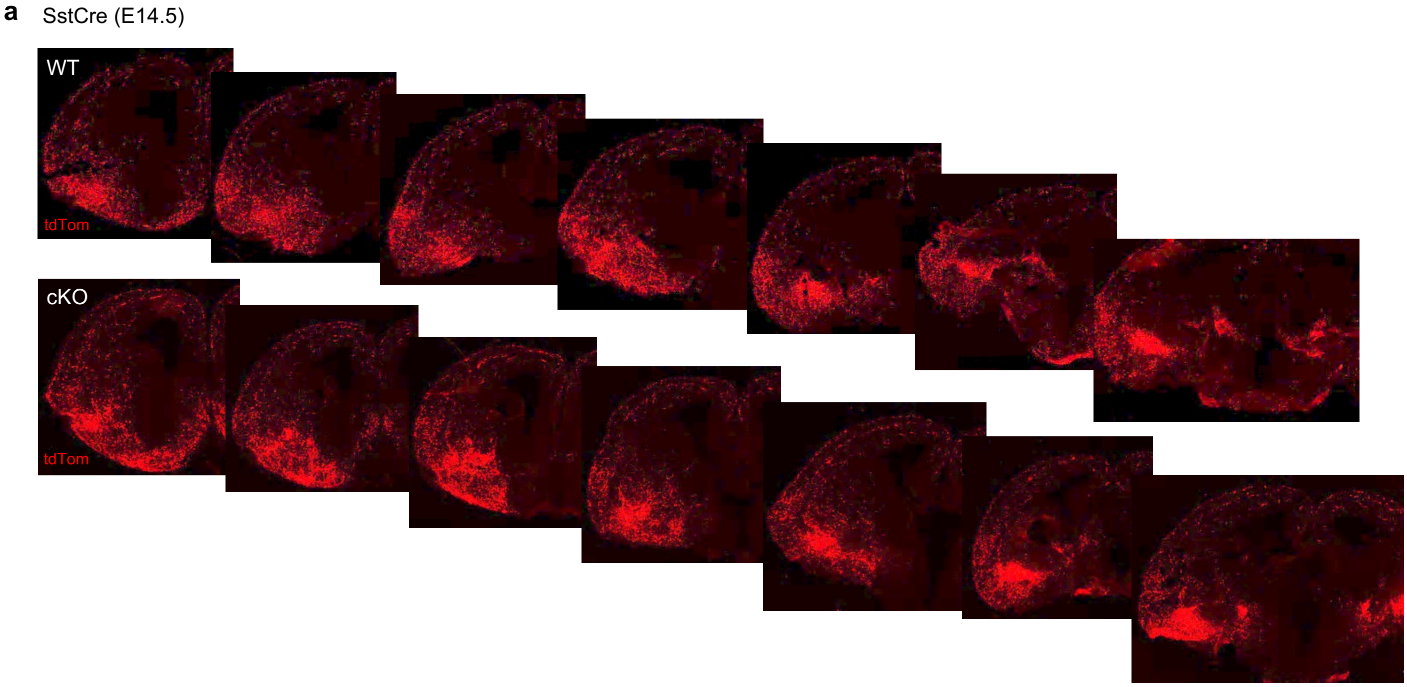

**Supplementary Fig 3**. **a** A representative series of coronal sections from *SstCreER-Arx* cKO or WT brains (E14.5) immunolabeled with tdTomato.

### Supplementary Figure 4

**a**

WT

Arx cKO (Gad2Cre)

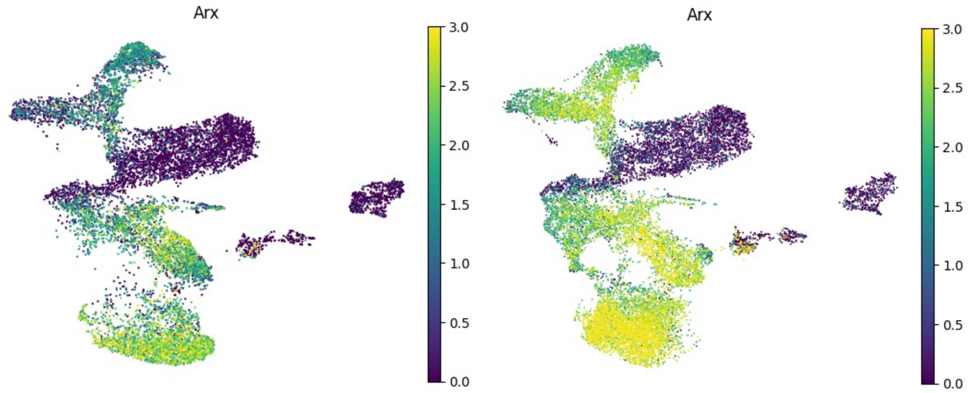


**b**

**Gene Regulatory network analysis by CellOracle**

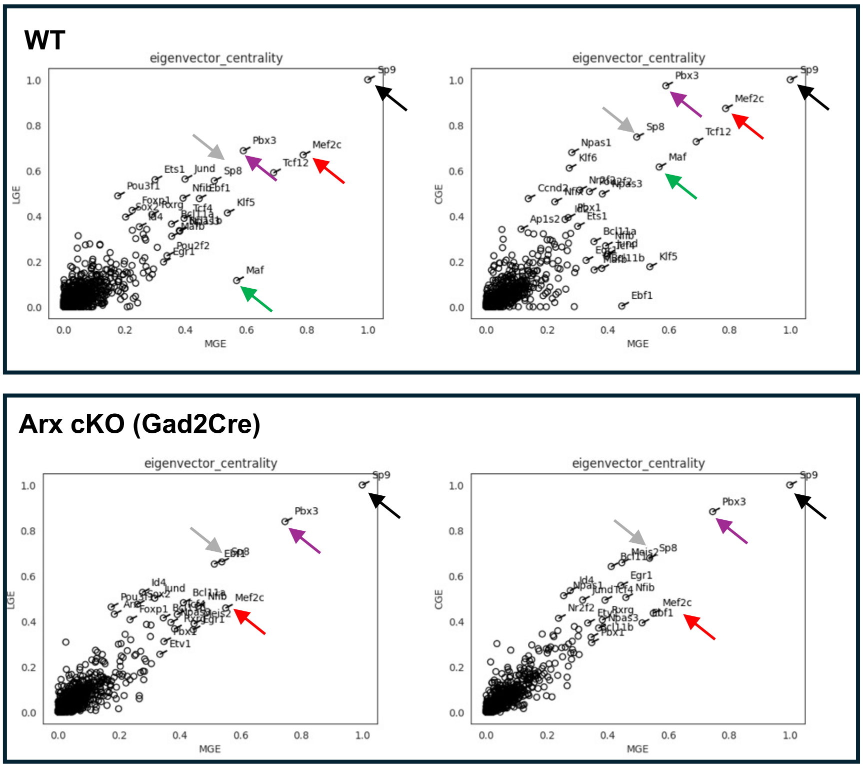

**Supplementary Fig 4**. **a** UMAP plot showing expression of *Arx* transcripts in *Gad2Cre-Arx* cKO and WT cell clusters. **b** Gene regulatory network analysis of *Gad2Cre-Arx* cKO and WT clusters using CellOracle. Color coded arrows indicate representative node genes in each cluster. Eigenvector centrality is a measure of the influence of a node in a network.

### Supplementary Figure 5

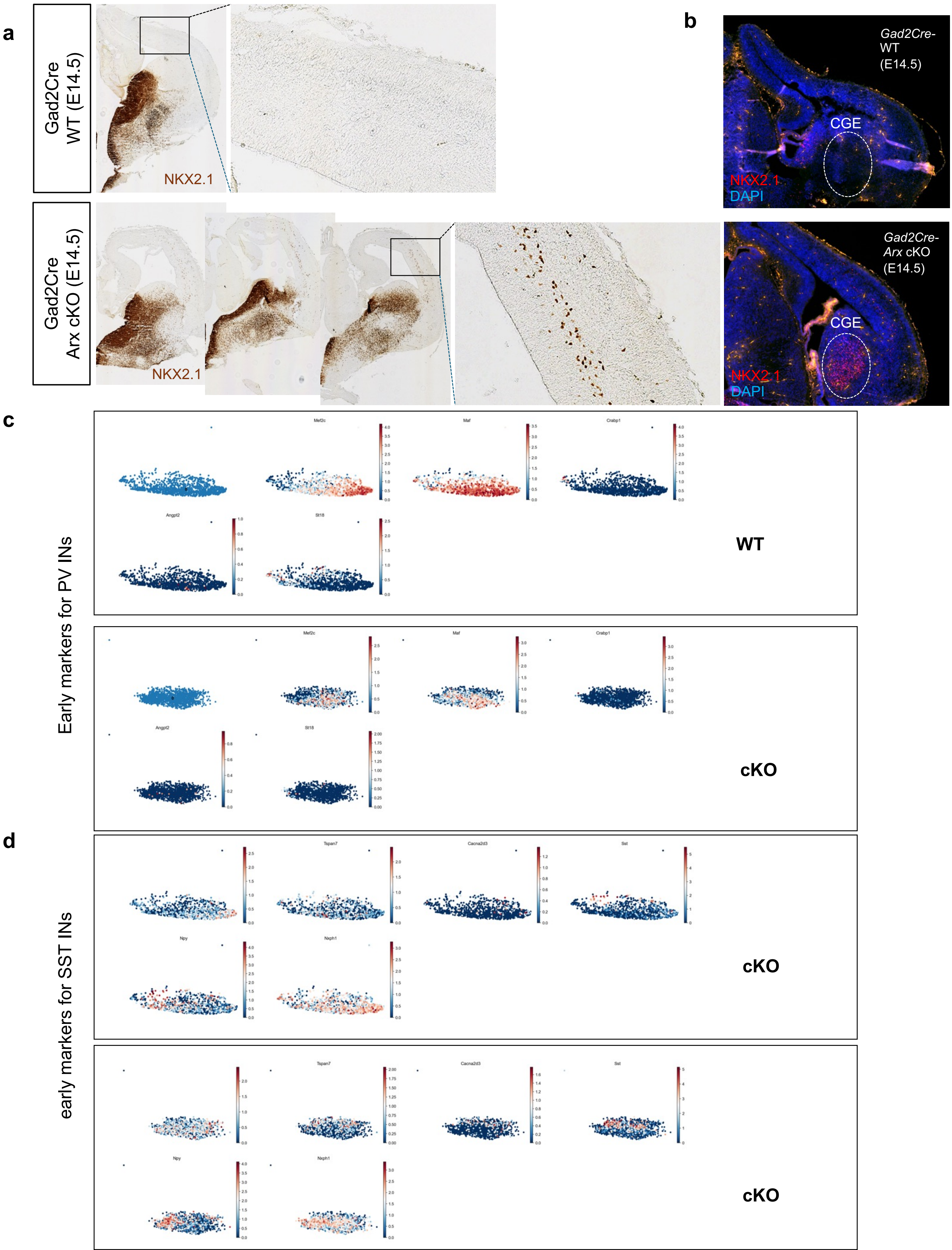

**Supplementary Fig 5**. **a, b** Representative images of the coronal sections of the *Gad2Cre-Arx* cKO and WT brains (E14.5) with NKX2.1 immunofluorescent labeling (a) or DAB-based immunohistochemical labeling (b). **c, d** UMAP plots of MGE clusters showing expression of early markers of PV (c) or SST fate (d) in *Gad2Cre-Arx* cKO and WT samples.

### Supplementary Figure 6

UMAP of Gad2 lineage cINs (ligands and receptors)

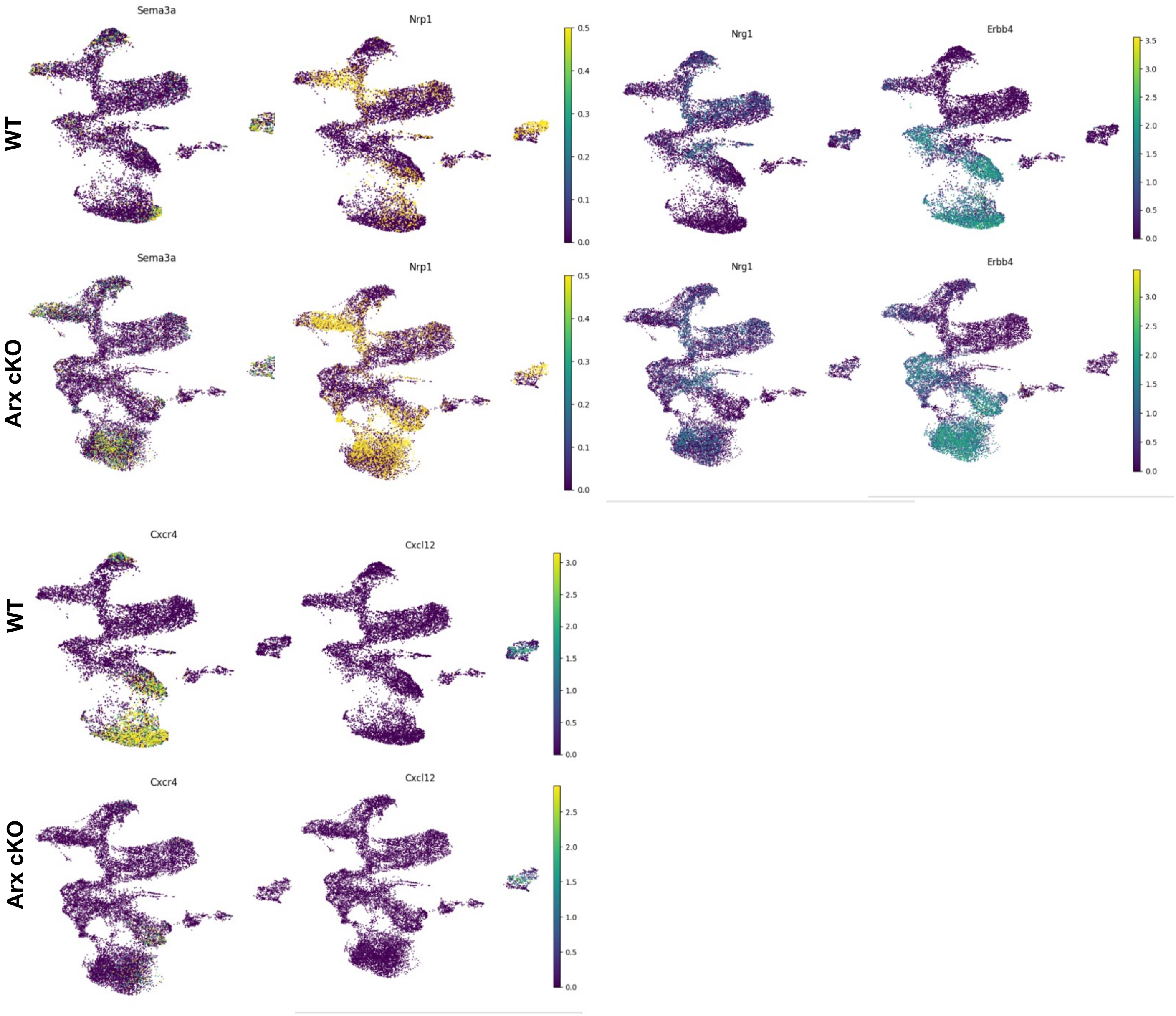

**Supplementary Fig 6**. **a** UMAP plot showing expression of ligands and receptors involved in IN migration cell migration in *Gad2Cre*-*Arx* cKO and WT cell clusters identified in scRNA-seq analysis.

### Supplementary Figure 7

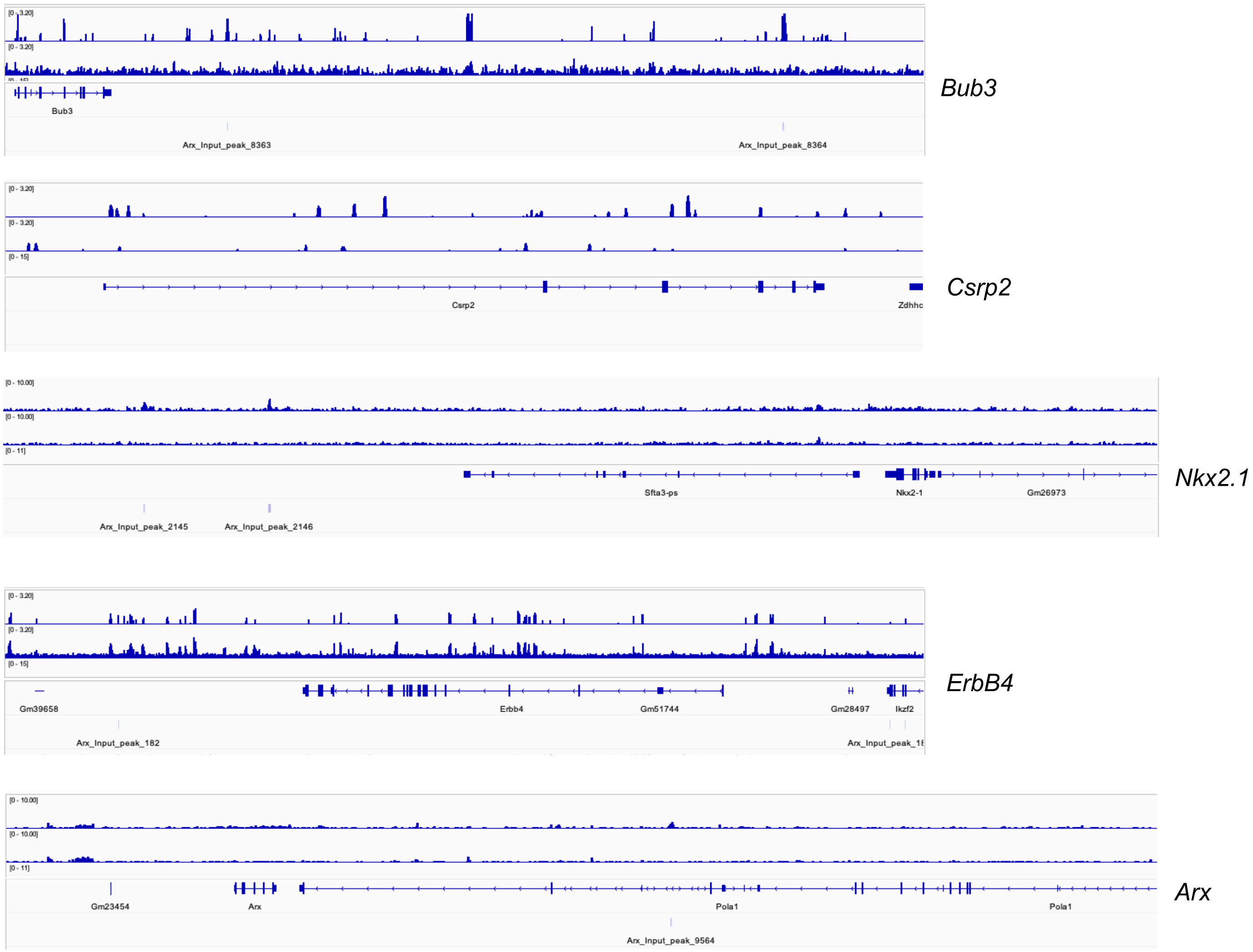

**Supplementary Fig 7**. **a** Examples of gene loci identified as ARX binding sites in ARX ChIP-seq analysis.

### Supplementary Figure 8

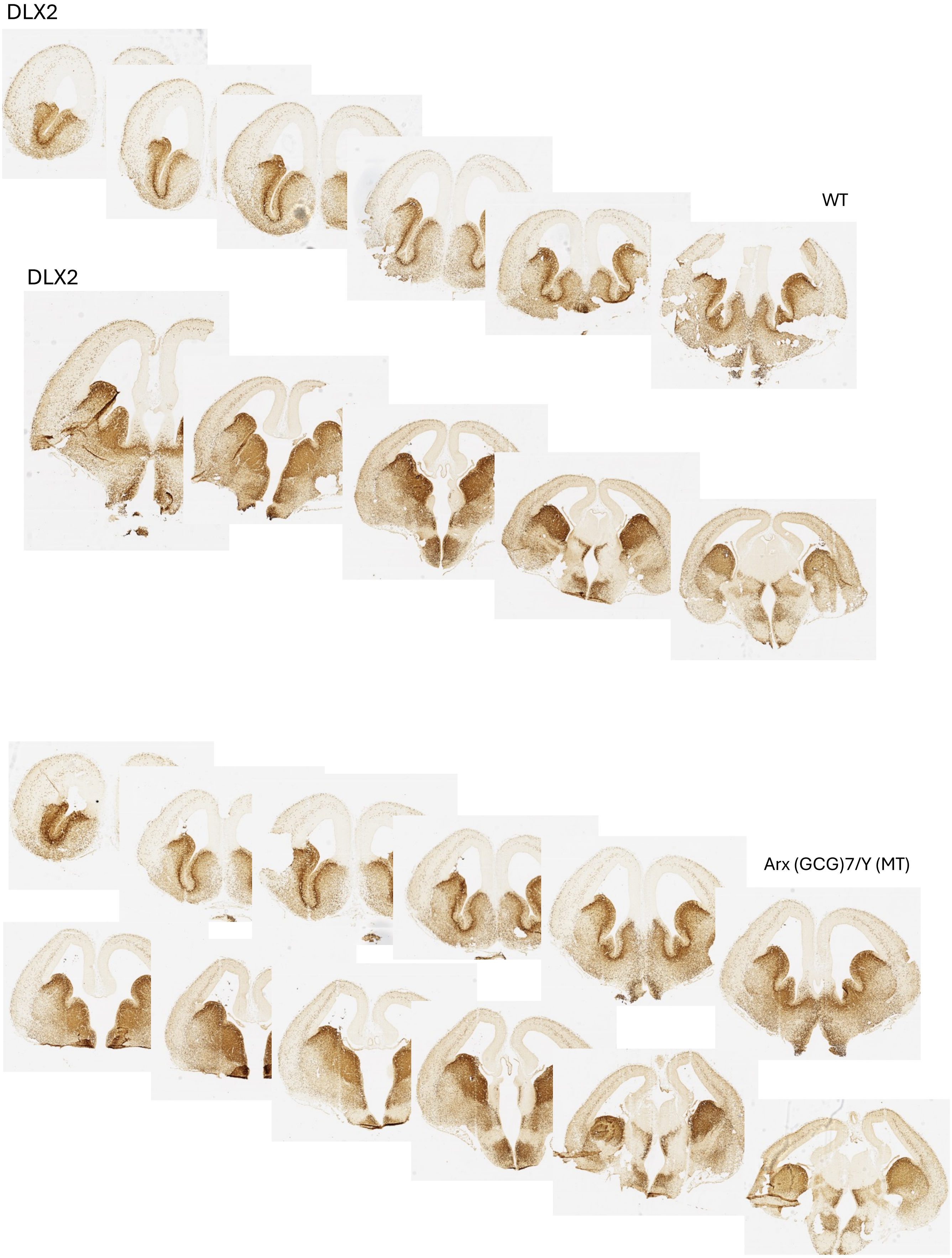

**Supplementary Fig 8**. **a** Representative series of images of DLX2 immunohistochemical labeling (DAB based) on coronal sections of the Arx (GCG)7 or WT brains.

### Supplementary Figure 9

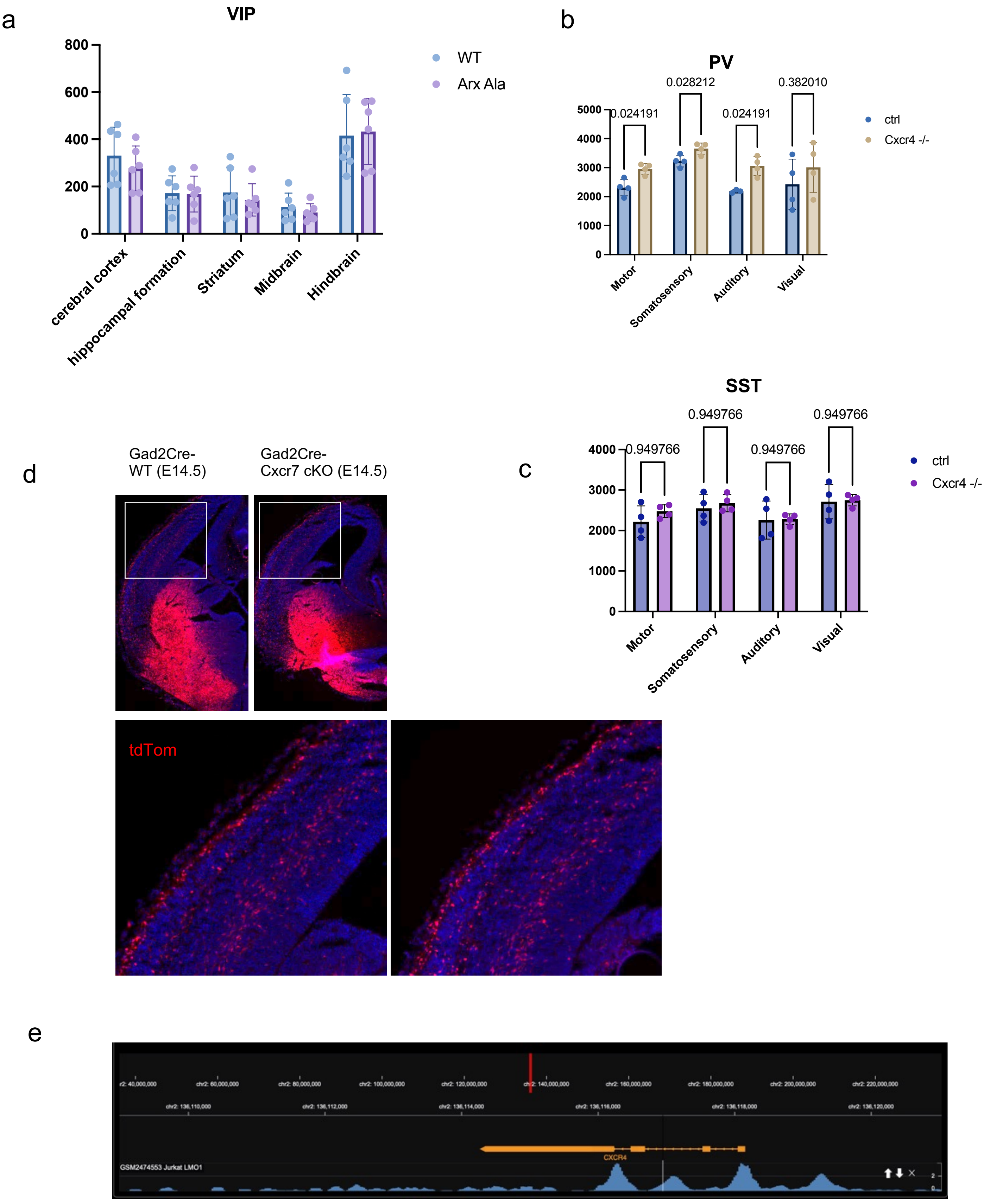

**Supplementary Fig 9**. **a** Quantification of VIP IN cell density (cells/mm^3) in each indicated region of the *Arx*$^{(GCG)7/Y}$ or WT brains, processed for 3D immunolabeling with VIP antibody after tissue clearing procedure. **b, c** Quantification of PV or SST IN cell density (cells/mm$^3$) in each indicated cortical layers of the *Gad2Cre*-*Cxcr4* cKO or WT brains, processed for 3D immunolabeling with VIP antibody after tissue clearing procedure. **d** Representative images of the tdTomato immunolabeling on coronal sections of the *Gad2Cre*-*Cxcr7* WT and cKO brains (E14.5). **e** LMO1 ChIP-seq peak analysis showing specific LMO1 peak at *Cxcr4* locus (ref. [82]).
